# Multi-omic analysis identifies mitochondrial dysfunction as a conserved driver of acute severity and long-term complications in RSV infection

**DOI:** 10.64898/2026.05.04.722656

**Authors:** Joseph Guarnieri, Nidia S. Trovao, Robert Schwartz

**Affiliations:** Guarnieri Research Group LLC, Philadelphia, PA 19130, USA; Blue Marble Space Institute of Science, Seattle, WA 98104, USA; Department of Pathobiology, College of Veterinary Medicine, University of Illinois Urbana-Champaign, Urbana, IL, USA; Carl R. Woese Institute for Genomic Biology, University of Illinois Urbana-Champaign, Urbana, IL, USA; National Center for Supercomputing Applications, University of Illinois; Weill Cornell Medicine, New York, NY 10065, USA

**Keywords:** Respiratory syncytial virus, mitochondrial dysfunction, oxidative phosphorylation, metabolic reprogramming, HIF-1α, post-viral complications

## Abstract

Respiratory syncytial virus (RSV) is a leading cause of lower respiratory tract infection in infants, older adults, and immunocompromised individuals. The molecular mechanisms linking acute RSV infection to disease severity and long-term complications remain incompletely understood.

Herein, we conducted a comprehensive multi-omic analysis of 12 independent datasets encompassing epigenomics, transcriptomics, proteomics, and metabolomics across diverse systems, including *in vitro* infection models, clinical cohorts, longitudinal pediatric studies, vaccination models, and multiple viral strains. Across these experimental platforms and omic analysis, RSV consistently triggered suppression of oxidative phosphorylation (OXPHOS), alongside HIF-1α-driven glycolytic metabolism and mitochondrial stress response. This coordinated reprogramming was consistent across transcriptomic, proteomic, and chromatin datasets.

In adult challenge studies, symptomatic individuals exhibited prolonged OXPHOS suppression and greater activation of HIF-1α immune signaling than asymptomatic individuals. Similarly, pediatric intensive care cohorts showed comparable signatures associated with severe disease. Vaccinated mice showed attenuation of infection-induced metabolic disruption, further supporting a link between mitochondrial dysfunction and disease severity. Longitudinal analyses in pediatric samples revealed that these metabolic alterations persist for up to 1-year post-infection, with sustained metabolic dysfunction, persistent epigenetic remodeling, and single-cell evidence of epithelial remodeling, including depletion of multiciliated cells, expansion of secretory populations, and prolonged OXPHOS suppression, in children who developed wheezing. Comparative analysis across RSV strains revealed variable OXPHOS suppression and variable HIF-1α activation, indicating strain-specific differences in metabolic reprogramming.

Together, these findings establish mitochondrial dysfunction as a central and conserved feature of RSV pathogenesis, encompassing acute severity, viral strain variation, and long-term complications, and highlight mitochondrial pathways as promising therapeutic targets to mitigate both acute disease severity and post-viral sequelae. Ultimately, demonstrating that distinct viral lineages drive unique bioenergetic phenotypes establishes a foundation for predictive molecular epidemiology and gaining insight into host-pathogen dynamics in response to novel interventions.

**Highlights:**

- Multi-omic integration of 13 independent RSV datasets reveals mitochondrial dysfunction as a conserved hallmark of infection.
- RSV consistently suppresses oxidative phosphorylation (OXPHOS) while activating HIF-1α signaling, glycolysis, mitochondrial stress responses, and immune pathways.
- Greater mitochondrial dysfunction correlates with increased disease severity, persists in children who develop wheezing, and is partially ameliorated by vaccination.
- Distinct RSV strains display variable patterns of metabolic reprogramming, linking viral genetic diversity to differential host mitochondrial responses.

## INTRODUCTION

Respiratory syncytial virus (RSV) is a leading cause of lower respiratory tract infection in infants, older adults, and immunocompromised populations worldwide, contributing to substantial morbidity, hospitalization, and long-term respiratory complications including recurrent wheezing and asthma^1^. Despite its major clinical burden, the host metabolic mechanisms that contribute to RSV severity and long-term sequelae remain incompletely understood.

Mitochondria are central regulators of cellular energetics, antiviral immunity, and oxidative stress responses, and growing evidence suggests they are frequent targets of respiratory viral infection^2–5^. We and others have previously identified mitochondrial dysfunction as a conserved feature of severe respiratory viral infections, including influenza A virus (IAV)^3^ and SARS-CoV-2, where viral infection induces rapid host bioenergetic reprogramming characterized by suppression of mitochondrial oxidative phosphorylation (OXPHOS), increased glycolytic reliance, and elevated mitochondrial reactive oxygen species (mROS)^2,6–9^. In SARS-CoV-2, viral proteins directly disrupt mitochondrial machinery^7,10–12^, repress mitochondrial transcripts^6,8,11^, and promote hypoxia-inducible factor 1 alpha (HIF-1α)-associated metabolic reprogramming^6,8,13,14^, that supports viral replication while amplifying inflammatory signaling through mitochondrial damage-associated molecular pattern release^2,7–9,15,16^. Notably, mitochondrial-targeted interventions have reduced respiratory viral replication and disease severity in experimental models^5,7,8,14^, suggesting mitochondrial dysfunction is a mechanistic contributor to pathogenesis rather than simply a downstream consequence of infection.

Accumulating evidence suggests RSV may induce similar mitochondrial perturbations. Prior studies have consistently demonstrated impaired mitochondrial respiration, decreased mitochondrial membrane potential (ΔΨm), and disruptions in mitochondrial dynamics^5,17,18^. Following RSV infection, selective degradation of nuclear factor erythroid 2-related factor 2 (NRF2) leads to increased levels of ROS^19–22^. RSV has been shown to stabilize HIF-1α and elevate HIF-1α target genes^23,24^. Treatment with the selective ROS^25,26^ and HIF-1α inhibitor^23^ decreases RSV replication. However, these findings remain fragmented across individual experimental systems, time points, and patient cohorts, and it remains unclear whether mitochondrial dysfunction represents a conserved feature of RSV infection across disease severity, viral strains, and long-term outcomes. In particular, whether these metabolic alterations persist beyond acute infection and contribute to post-RSV complications remains poorly defined.

The global circulation of RSV is characterized by the continuous replacement of dominant lineages, driven by genetic drift and episodic selective sweeps^27^. Genomic epidemiology has demonstrated that structural variations, such as the 72-nucleotide duplication in the G gene of the RSV-A ON1 lineage and the 60-nucleotide duplication in the RSV-B BA lineage, facilitate rapid global dissemination and localized epidemic waves^28^. However, the extent to which these macro-evolutionary genomic shifts dictate micro-level host-pathogen interactions, specifically concerning the host bioenergetic collapse, remains poorly characterized. Bridging this gap requires evaluating whether contemporary viral lineages exert distinct metabolic pressures on the host compared to historic strains.

Here, we performed a comprehensive multi-omic analysis of RSV-associated mitochondrial dysfunction by integrating 13 independent datasets spanning transcriptomics^29–35^, epigenomics^30,36^, proteomics^37,38^, metabolomics^39,40^, and single-nucleus RNA-sequencing (snRNA-seq)^29^ across *in vitro* models, acute patient cohorts, longitudinal pediatric samples, and vaccination studies. This systems-level approach enabled us to define conserved and context-specific metabolic responses across infection stages, disease severity, viral strains, and long-term outcomes.

Across datasets, we identified a conserved RSV-associated metabolic signature characterized by suppression of OXPHOS, activation of HIF-1α-associated glycolytic pathways, and induction of mitochondrial stress responses across epigenetic, transcriptional, proteomic, and metabolomic layers. These alterations were amplified in severe disease, persisted in pediatric cohorts with long-term wheezing complications, varied across viral strains, and were partially attenuated by vaccination. Collectively, our findings identify mitochondrial dysfunction as a conserved feature of RSV pathogenesis and suggest that host mitochondrial metabolism may represent a therapeutic target for both acute disease and post-viral complications.

## RESULTS

### Multi-Omic integration of RSV infection reveals targeted downregulation of OXPHOS transcripts alongside HIF-1**α** and mitochondrial stress response activation

#### RNA-seq, ATAC-seq, and Proteomics Analyses of In Vitro Infections Across Human Airway Models

To investigate the molecular mechanisms by which RSV infection alters mitochondrial and immune responses, we performed a meta-analysis of published multi-omic datasets^30–33,38^. Studies were curated and stratified by time post-infection, cell type, and omics modality. We then applied our custom pathway^6,9^ analysis pipeline to reanalyze transcriptomic and proteomic datasets (ATAC-seq, RNA-seq, and proteomics). Detailed experimental information for each dataset is provided in **Supplemental Table 1**. Gene set enrichment analysis (GESA) was performed to calculate normalized enrichment scores (NES), with significance defined as FDR < 0.25, and results visualized using a heatmap (**Fig. 1**). Strikingly, we observed consistent suppression of mitochondrial OXPHOS gene expression alongside robust upregulation of HIF-1α signaling across studies. These changes were accompanied by enrichment of immune activation and cell death pathways, as well as mitochondrial stress responses, including pathways associated with mitochondrial DNA (mtDNA) and double-stranded RNA (mtdsRNA) release.

**Figure 1.**
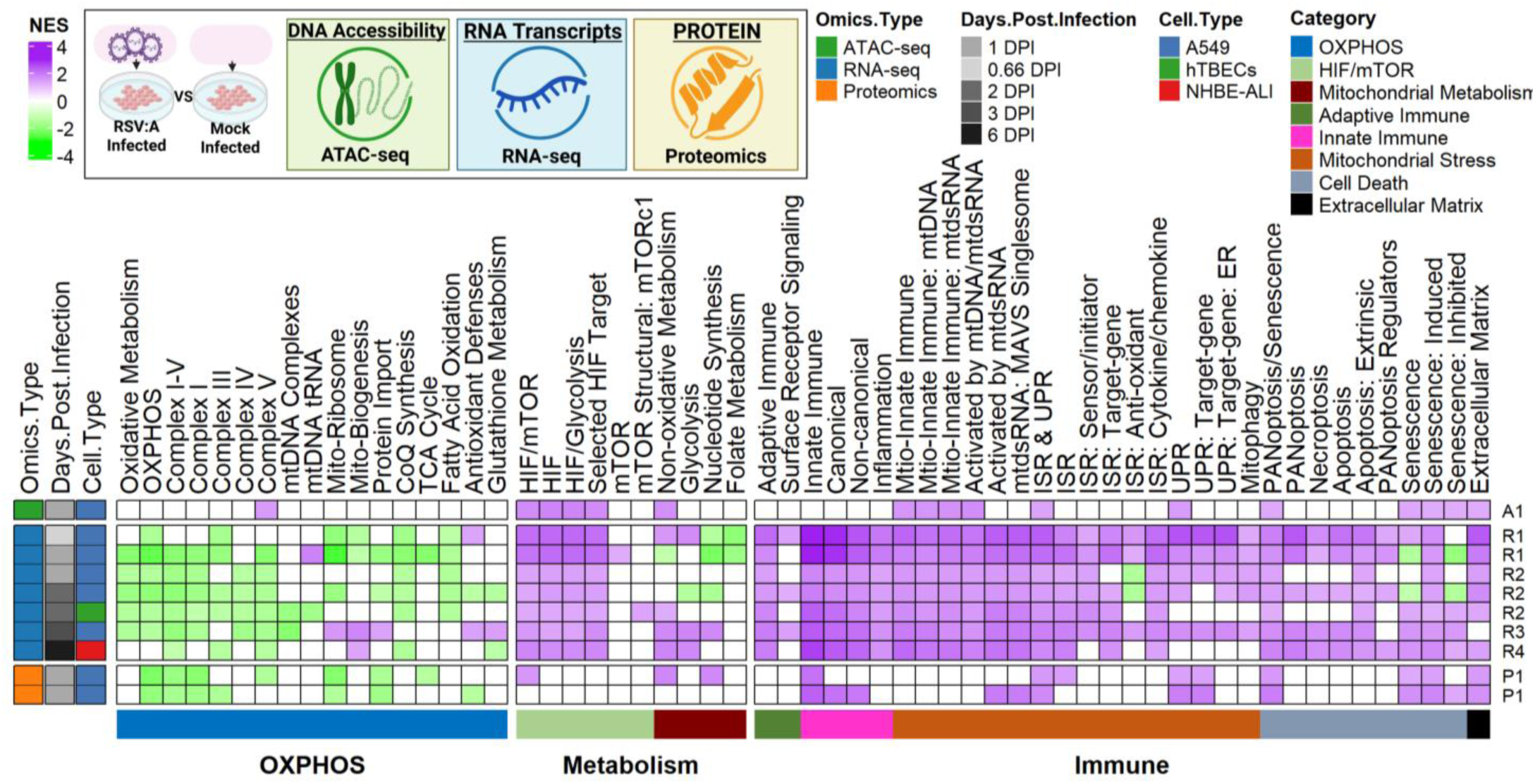
RSV induces conserved suppression of OXPHOS and activation of HIF-1α signaling across *in vitro* airway models. (A) Schematic overview of multi-omics datasets analyzed. Heatmap displays key metabolic and immune pathways identified by gene set enrichment analysis (GSEA; normalized enrichment score [NES], false discovery rate [FDR] < 0.25) comparing RSV A2-infected versus mock-infected samples, shown across omics platforms, days post-infection (DPI), and cell types. Key metabolic and immune pathways were curated using custom gene sets as previously described (^6,9^). Data analyzed from each study is designated by the right annotation: A1(^28^); R1(^28^); R2(^29^); R3(^29^); R4(^31^); P1(^36^).

At 1 day-post-infection (DPI), ATAC-seq analysis revealed increased chromatin accessibility at loci associated with HIF-1α and mitochondrial stress response genes, indicating rapid epigenetic reprogramming. Consistent with these findings, transcriptomic analyses at 0.75 and 1 DPI demonstrated significant upregulation of HIF-1α-associated pathways, mitochondrial stress responses, and immune signaling programs. Concomitantly, there was marked downregulation of genes involved in oxidative metabolism, including OXPHOS, the tricarboxylic acid (TCA) cycle, and fatty acid oxidation. To validate whether these transcriptional changes were reflected at the protein level, we analyzed proteomic datasets at 1 DPI. Consistent with transcriptomic findings, proteomic analysis demonstrated downregulation of OXPHOS components alongside upregulation of HIF-1α signaling, mitochondrial stress pathways, and immune responses, confirming coordinated regulation across molecular layers (**Fig. 1**). Together these combined Omic changes, characterized by OXPHOS suppression, HIF-1α activation, and induction of mitochondrial stress and immune pathways, were conserved across cell type analyzed and collection point, indicating that they represent a conserved host response to RSV infection.

#### RNA-seq and snRNA-seq Analyses of RSV Infection in Human Nasal Samples

To extend our findings into clinically relevant populations, we analyzed nasal curettage samples from healthy adult volunteers (18-55 years) following natural RSV infection. Samples were collected at <1, 3, and 7 DPI and stratified into symptomatic and asymptomatic groups^34,35^ (**Fig. 2**). Consistent with our *in vitro* observations, transcriptional profiling in adult cohorts revealed suppression of OXPHOS alongside upregulation of HIF-1α signaling, mitochondrial stress responses, and immune pathway activation across all timepoints, further validating the importance of these mechanisms in RSV pathogenesis. Notably, at 7 DPI, symptomatic patients exhibited more pronounced downregulation of OXPHOS and more robust upregulation of HIF-1α and immune pathways relative to asymptomatic individuals, suggesting a relationship between transcriptional dysregulation and symptom severity.

**Figure 2.**
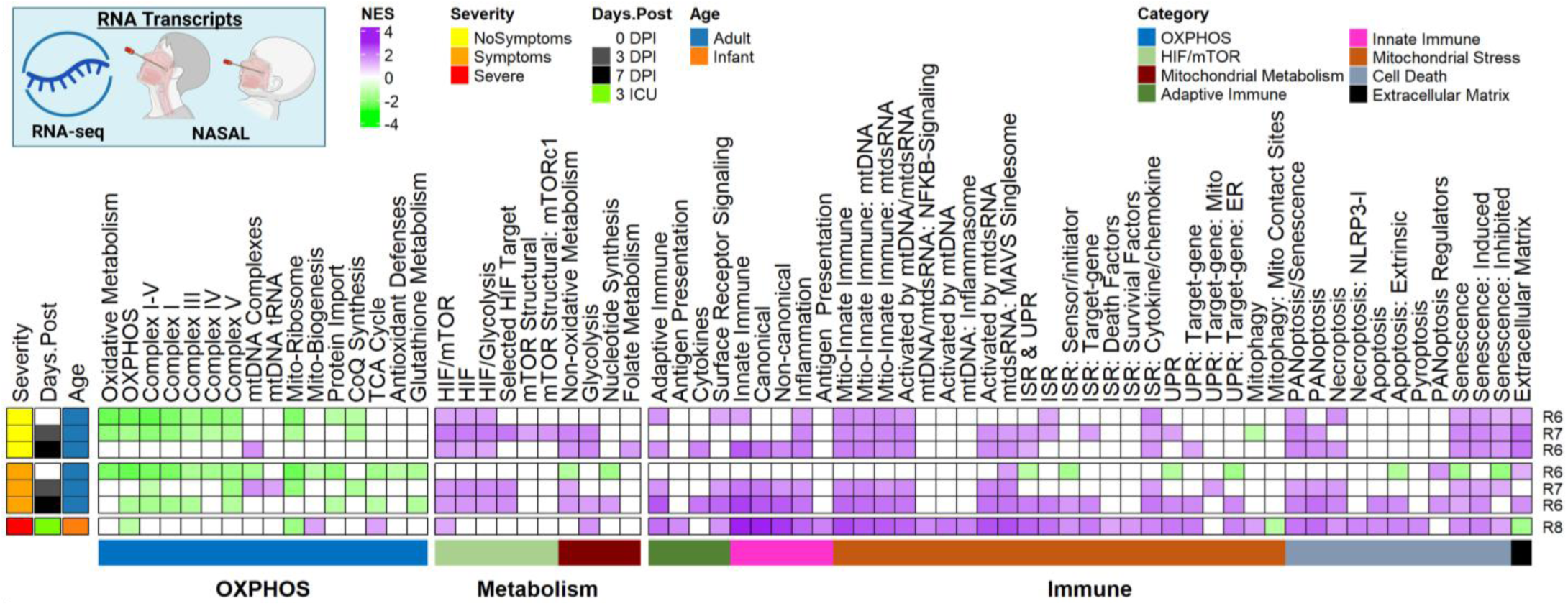
Mitochondrial dysfunction tracks with symptom severity in adult and pediatric RSV cohorts. Heatmap of significantly enriched metabolic and immune pathways (GSEA; NES, FDR < 0.25) derived from RNA-seq data of nasal samples from RSV-infected adult and pediatric cohorts. Nasal curettage samples from healthy adult volunteers (18-55 years) collected at <1, 3, and 7 DPI, comparing symptomatic (RSV+Symptoms+) and asymptomatic (RSV+Symptoms−) individuals to RSV-negative controls (RSV−). Nasopharyngeal samples from pediatric subjects (<2 years) collected at 3 days post-ICU (DP-ICU) admission, comparing severe to mild RSV cases. Key metabolic and immune pathways were curated using custom gene sets as previously described (^6,9^). Data analyzed from each study is designated by the right annotation: R6(^32^); R7(^33^); R8(^33^).

To investigate whether these patterns extended to more severe disease, we analyzed nasopharyngeal samples from pediatric subjects (<2 years) collected at 3 days post-ICU (DP-ICU) admission stratified between severe and mild conditions^35^ (**Fig. 2**). Compared with mild patients, severe RSV patients again exhibited sustained suppression of OXPHOS alongside upregulation of HIF-1α and pro-inflammatory immune pathways, suggesting that OXPHOS impairment alongside HIF1a and immune pathways activation is associated with severity.

To further interrogate transcriptional differences associated with disease severity, we performed Gene Ontology (GO) enrichment analysis and volcano plot-based differential expression analysis (**Sup Fig. 1, 2**). In adults, comparison of symptomatic versus asymptomatic cases demonstrated significant enrichment of sustained inflammatory and immune response pathways in severe disease (**Sup Fig. 1A)**. In contrast, asymptomatic cases showed enrichment of multiciliated epithelial cell-associated pathways, suggesting epithelial recovery. These trends were similarly observed in severe infant cohorts (**Sup Fig. 2A)**. Volcano plot of differential expression genes revealed a greater magnitude of transcriptional changes in severe cases in both adults and infants compared to mild cases, consistent with more extensive transcriptional reprogramming in severe disease and more rapid resolution in milder infections (**Sup Fig. 1B, 2B**). Collectively, these findings indicate that OXPHOS impairment and upregulation of HIF-1α signaling, mitochondrial stress pathways, and immune responses are conserved features associated with RSV-severity. To further investigate whether metabolic and immune alterations are associated with RSV severity, we examined whether vaccination, which reduces disease severity, also mitigates the metabolic alterations associated with RSV infection.

#### RSV vaccination in Rodents attenuates infection-induced metabolic reprogramming

To determine whether these metabolic and immune alterations track with disease severity, we next assessed the impact of vaccination, a known modifier of RSV severity, on infection-induced metabolic reprogramming^40^ (**Fig. 3**). Serum metabolomic profiling of RSV-infected mice revealed that compared to vaccinated (primed) animals, unvaccinated animals exhibited signatures of mitochondrial dysfunction, including elevated lactate levels indicative of increased glycolysis and reduced mitochondrial oxidative metabolism, along with increased immunometabolites such as kynurenine and itaconate (**Fig. 3**). In contrast, vaccinated mice showed attenuation of these metabolic perturbations and preservation of metabolic homeostasis (**Fig. 3B**). Together, these findings support a link between RSV disease severity and mitochondrial dysfunction and demonstrate that vaccination mitigates infection-induced metabolic reprogramming.

**Figure 3.**
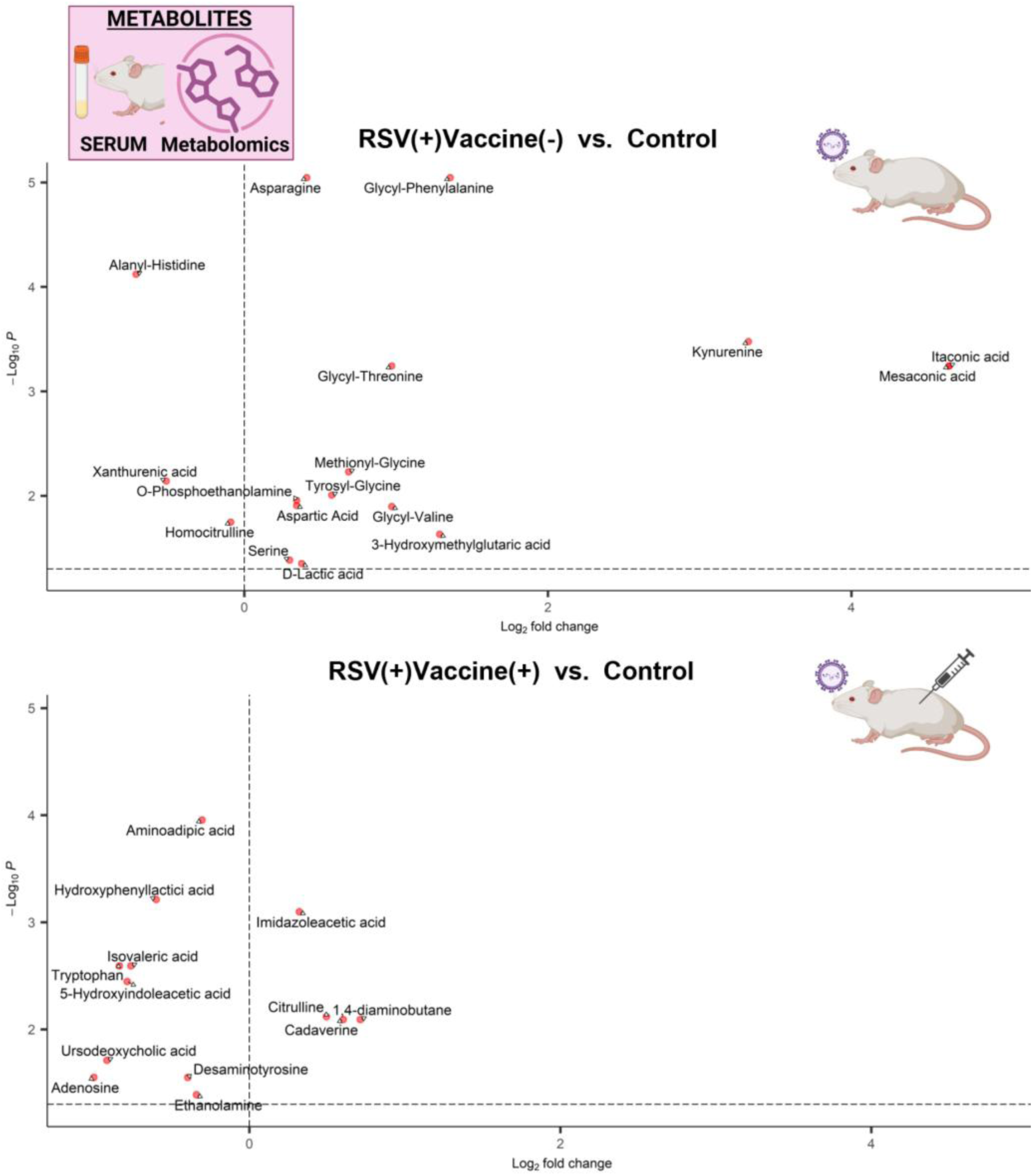
Vaccination attenuates RSV-induced metabolic reprogramming *in vivo*. Volcano plots of differentially abundant metabolites in serum from BALB/c mice at 7 DPI, comparing RSV-infected non-vaccinated (RSV+TriAdj−) (A) and RSV-infected vaccinated (RSV+TriAdj+) (B) mice to RSV-negative controls. Each point represents a metabolite, plotted by log_₂_ fold change (x-axis) versus −log_₁₀_ adjusted p-value (y-axis). Data was analyzed from M2(^38^).

### Persistent complications associated with RSV-infect are associated with prolonged metabolic reprogramming

Recent attention to post-viral syndromes, particularly in the context of COVID-19, has highlighted the persistence of mitochondrial dysfunction following viral clearance, especially in severe cases and individuals with post-acute sequelae^41,42^. Similarly, RSV has long been associated with post-acute complications, including chronic respiratory conditions such as wheezing and asthma in pediatric populations^1^. To determine whether mitochondrial dysfunction persists after RSV infection, we analyzed longitudinal infant-derived samples primary nasal airway epithelial cells (NAECs) collected ∼1-year post-infection (YPI)^29^, and blood^36^ and serum^39^ datasets collected 1.5-2.5 YPI.

#### scRNA-seq of RSV infection reveals loss of multiciliated cells and metabolic reprogramming across epithelial and immune populations

We first analyzed snRNA-seq data from differentiated primary NAEC cultures obtained from children aged 2-3 years enrolled in a longitudinal study that collected samples at 1 YPI, stratified by prior RSV infection status and wheeze phenotype^29^. Relative cell population abundances were calculated across three groups: non-infected non-wheezers (RSV−Wheeze−), RSV-infected non-wheezers (RSV+Wheeze−), and RSV-infected wheezers (RSV+Wheeze+) (**Fig. 4A**). Major cell populations identified included secretory cells, multiciliated epithelial cells, basal cells, and T cell lineages. Cell-type composition analysis revealed a significant enrichment of secretory cell populations and a concomitant reduction in multiciliated and basal epithelial cells following RSV infection (**Fig. 4B**). Notably, this shift was more pronounced in RSV-infected wheezers, suggesting a link between epithelial remodeling and disease severity. In parallel, RSV infection was associated with an expansion of T cell populations, consistent with enhanced immune activation.

**Figure 4.**
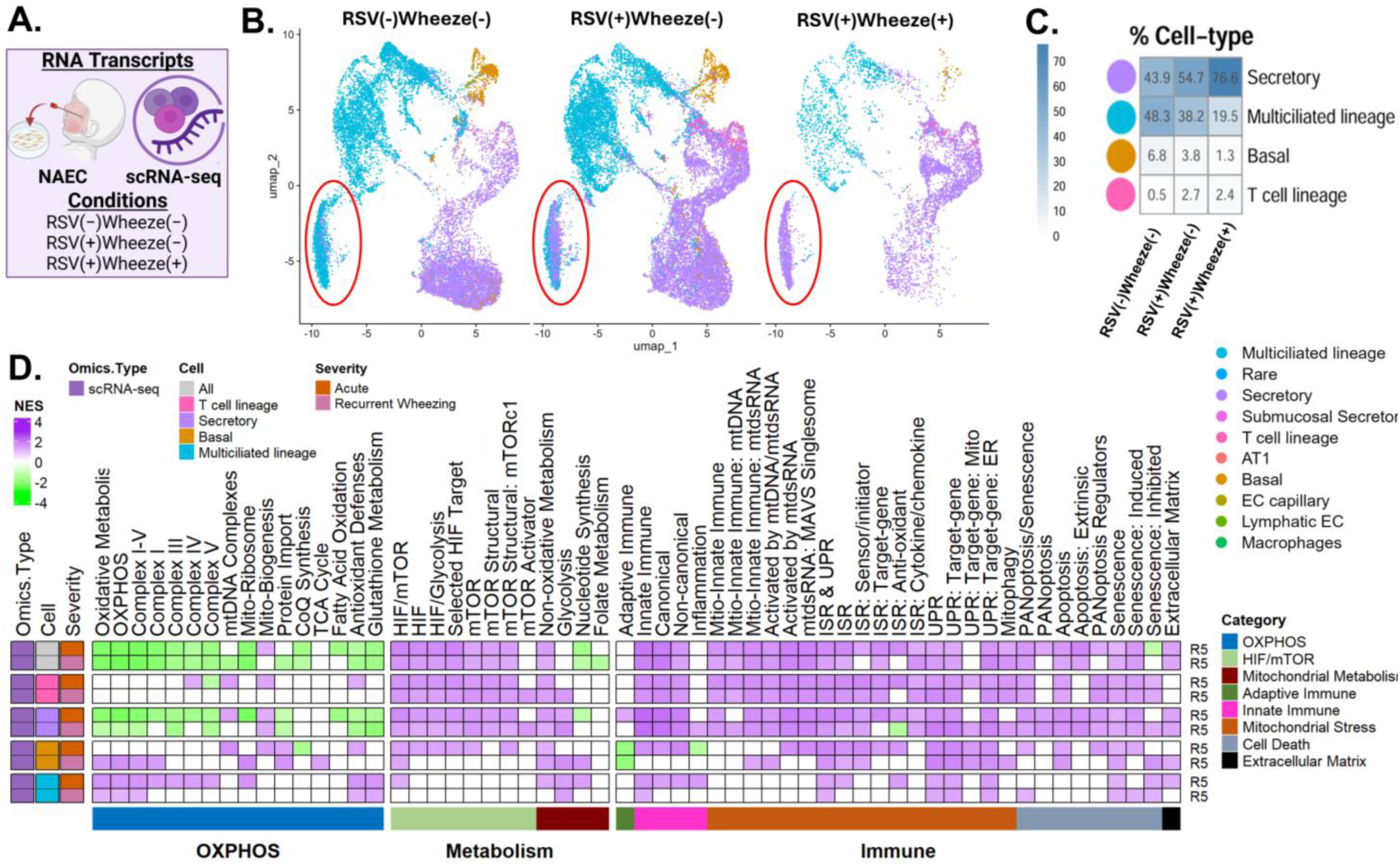
Single-cell analysis reveals persistent epithelial remodeling and metabolic dysfunction after RSV infection. (A) UMAP projection of single nuclei from differentiated primary NAEC cultures. (B) Heatmap showing relative cell-type abundances from differentiated primary NAEC cultures obtained from children (2-3 years) at 1 YPI, colored by annotated cell type across RSV−Wheeze−, RSV+Wheeze−, and RSV+Wheeze+ groups. (C) Heatmap of normalized enrichment scores (GSEA, NES) for significantly enriched pathways (FDR < 0.25) from comparative analyses of RSV+Wheeze+ and RSV+Wheeze− relative to RSV−Wheeze− controls. Key metabolic and immune pathways were curated using custom gene sets as previously described (^6,9^). Data from each study are indicated by the right annotation R5(^30^).

Next, we performed comparative analyses of RSV-infected wheezers and RSV-infected non-wheezers relative to non-infected non-wheezer controls (**Fig. 4C**). RSV infection was again associated with suppression of OXPHOS pathways alongside upregulation of HIF-1α signaling, mitochondrial stress responses, and immune pathways. At single-cell resolution, upregulation of HIF-1α and non-oxidative glycolytic and mitochondrial stress response pathways was evident across multiple cell types. Notably, multiciliate epithelial cells also displayed robust suppression of OXPHOS pathways, correlating with the observed depletion of this cell population. In contrast, T cell populations exhibited increased metabolic activity, including elevated expression of both OXPHOS and glycolytic pathways, consistent with activation-associated metabolic reprogramming (**Fig 4C**). Collectively, these findings demonstrate that RSV-infection induces persistent cell-type-specific metabolic remodeling characterized by loss of multiciliated epithelial cells and expansion of secretory and immune populations, particularly in disease states associated with wheezing.

#### Persistent metabolic and epigenetic alterations following RSV-infection suggest long-term mitochondrial dysfunction associated with post-viral sequelae

To determine whether metabolic and epigenetic alterations persist following RSV infection, we analyzed longitudinal infant cohorts 1 YPI, comparing individuals who developed wheezing to those who did not^36,39^. Serum metabolomic profiling revealed decreased levels of TCA cycle intermediates alongside increased lactate, consistent with a shift toward glycolytic metabolism and reduced mitochondrial oxidative function (**Fig 5A**). To further investigate regulatory mechanisms underlying these metabolic changes, we analyzed DNA methylation data using bisulfite sequencing (**Fig 5B**). This revealed decreased methylation at loci associated with HIF-1α signaling, mitochondrial stress response pathways, and cell fate programs including apoptosis and senescence, suggesting increased transcriptional accessibility of these pathways. Collectively, these findings support a model in which RSV infection induces persistent metabolic and mitochondrial dysfunction that extends beyond the acute phase of infection. These alterations may be exacerbated in individuals with underlying conditions and could contribute to long-term post-RSV complications, including wheezing and chronic respiratory disease.

**Figure 5.**
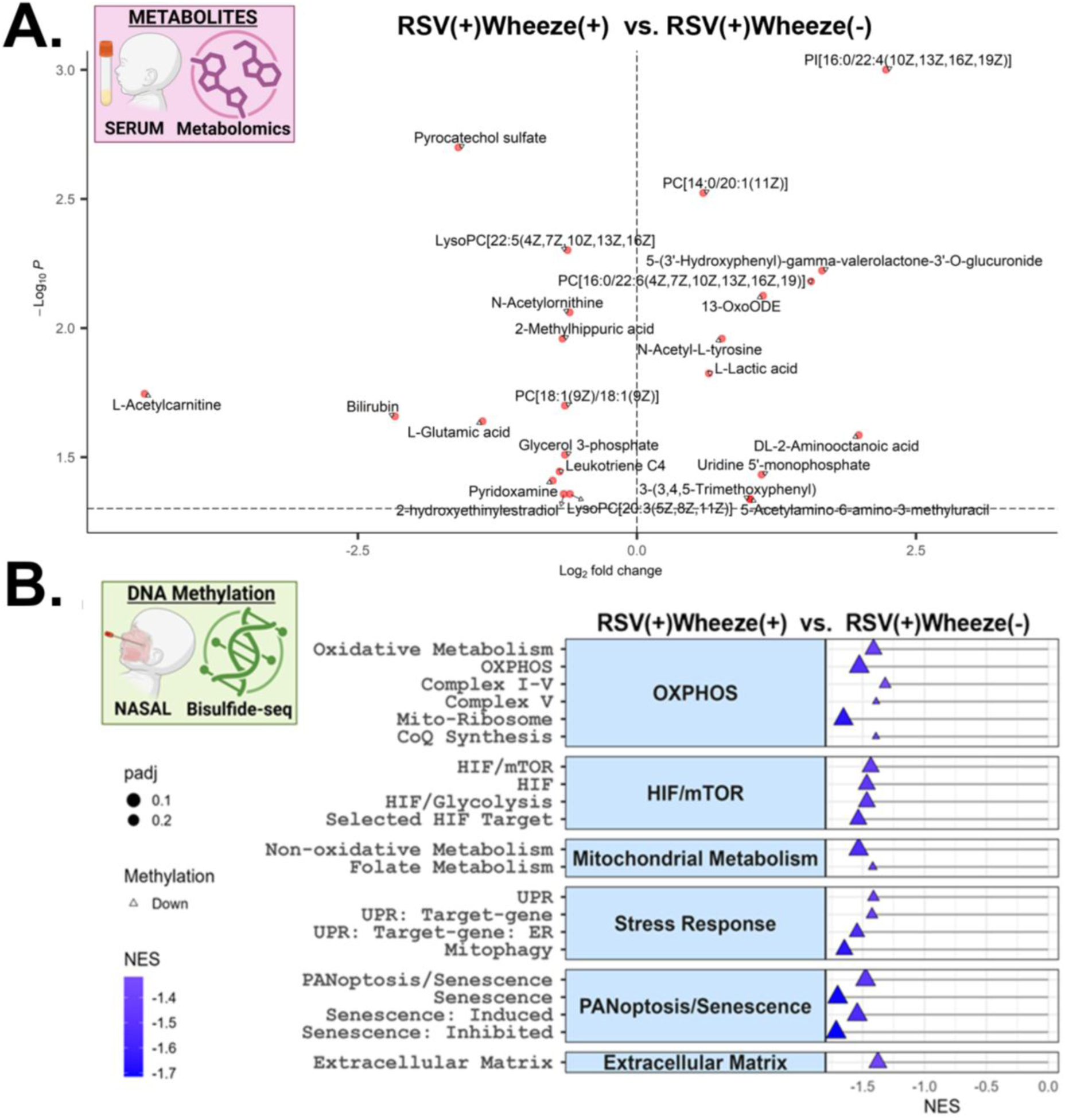
Persistent metabolic and epigenetic remodeling following RSV infection is associated with wheezing outcomes. (A) Volcano plot of significantly altered serum metabolites identified by metabolomic analysis in pediatric patients up to 1 YPI, comparing children with and without persistent RSV-associated wheezing. Each point represents a metabolite, plotted by log_₂_ fold change (x-axis) versus −log_₁₀_ adjusted p-value (y-axis). Data was analyzed from M1(^37^). (B) Lollipop plot displaying GSEA-based pathway enrichment (NES, FDR < 0.25) from DNA methylation (bisulfite sequencing) analysis of nasal samples collected up to one year post-infection, comparing children with and without persistent RSV-associated wheezing. Key metabolic and immune pathways were curated using custom gene sets as previously described (^6,9^). Data analyzed from: B1(^34^).

### RSV strain-specific differences in metabolic reprogramming highlight variable regulation of OXPHOS, HIF-1α, and mTOR signaling

Collectively, our multi-omic analyses across *in vitro* systems and patient samples revealed a consistent metabolic signature of RSV infection characterized by suppression of OXPHOS, activation of HIF-1α signaling, and induction of mitochondrial stress and immune response pathways. To determine whether these effects are strain-specific, we compared transcriptional responses in A549 cells infected with two RSV-A strains (A2 and ON1) and two RSV-B strains (B:18537 and BA)^32^ (**Fig 6A**). While OXPHOS suppression was observed across all strains, its magnitude was reduced in cells infected with RSV B:18537 relative to the other strains (**Fig 6B**). Notably, RSV B:18537 uniquely exhibited enrichment of mTOR signaling, whereas the remaining strains showed consistent upregulation of HIF-1α pathways, in agreement with our prior analyses. Despite these metabolic differences, all strains induced broadly comparable activation of immune-related pathways based on gene set enrichment analysis, suggesting that the primary strain-specific variation lies in metabolic reprogramming rather than immune activation.

**Figure 6.**
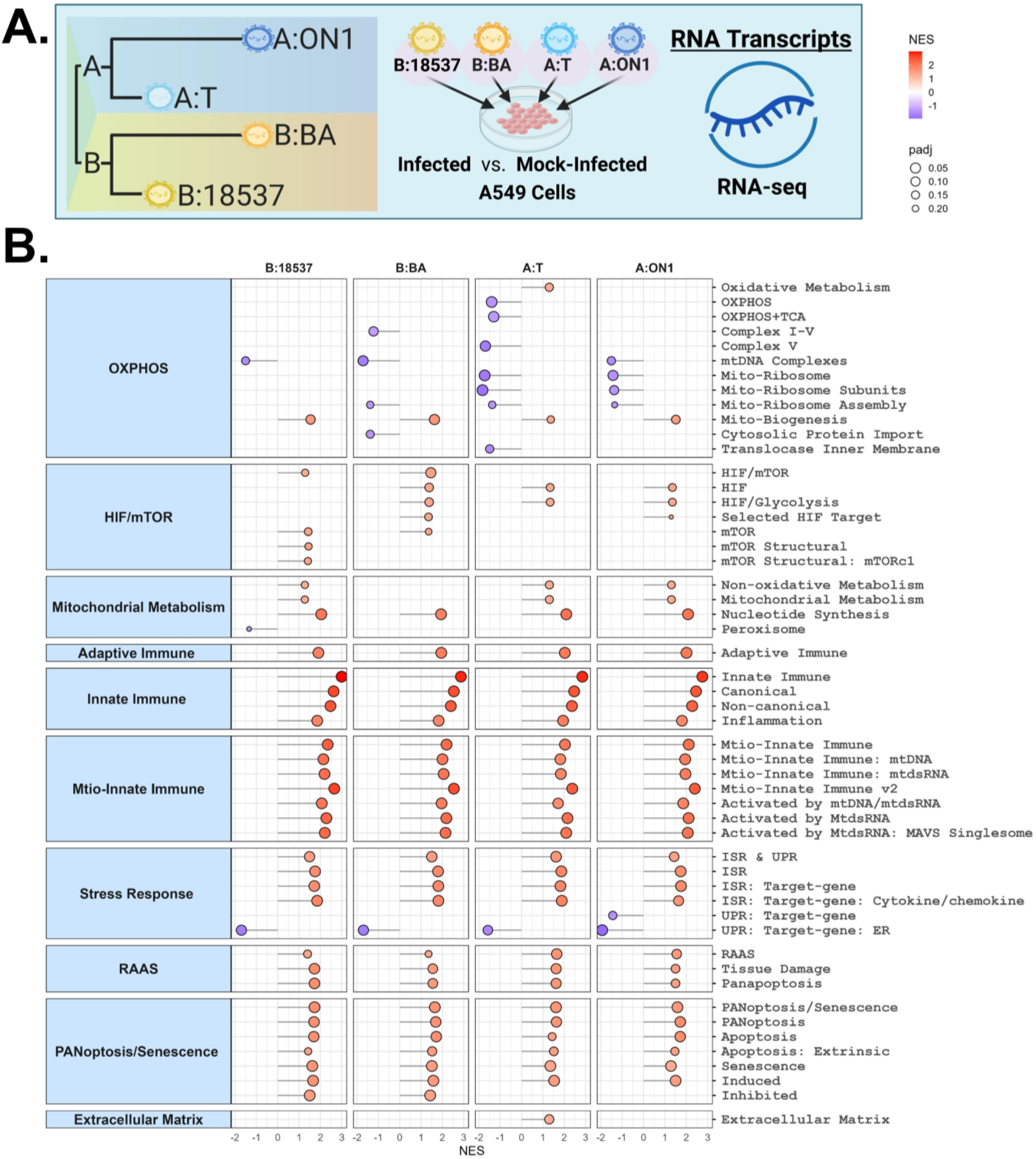
RSV strains exhibit distinct metabolic reprogramming. (A) Phylogenetic tree depicting the RSV strains analyzed by RNA-seq. (B) Lollipop plots showing lineage-specific alterations detected by GSEA (NES; FDR < 0.25). Key metabolic and immune pathways were curated using custom gene sets as previously described (^6,9^). Data analyzed from: (R9(^30^)).

GO enrichment and volcano plot analyses further supported these findings, with RSV B:18537 displaying the least extensive transcriptional and pathway-level perturbations among the strains examined (**Sup Fig 3A-B**). Taken together, these results demonstrate that emergent RSV strains exhibit distinct metabolic signatures, particularly in the regulation of OXPHOS and HIF-1α pathways.

Collectively, these multi-omic analyses across *in vitro* systems, acute clinical cohorts, and longitudinal patient samples identify a conserved RSV-induced metabolic program characterized by suppression of OXPHOS and activation of HIF-1α-driven mitochondrial stress and immune pathways (**Summarized in Fig 7**). The consistency of this signature across disease severity, persistence after infection, and attenuation with vaccination supports a central role for mitochondrial dysfunction in RSV pathogenesis and its associated short-and long-term complications.

**Figure 7.**
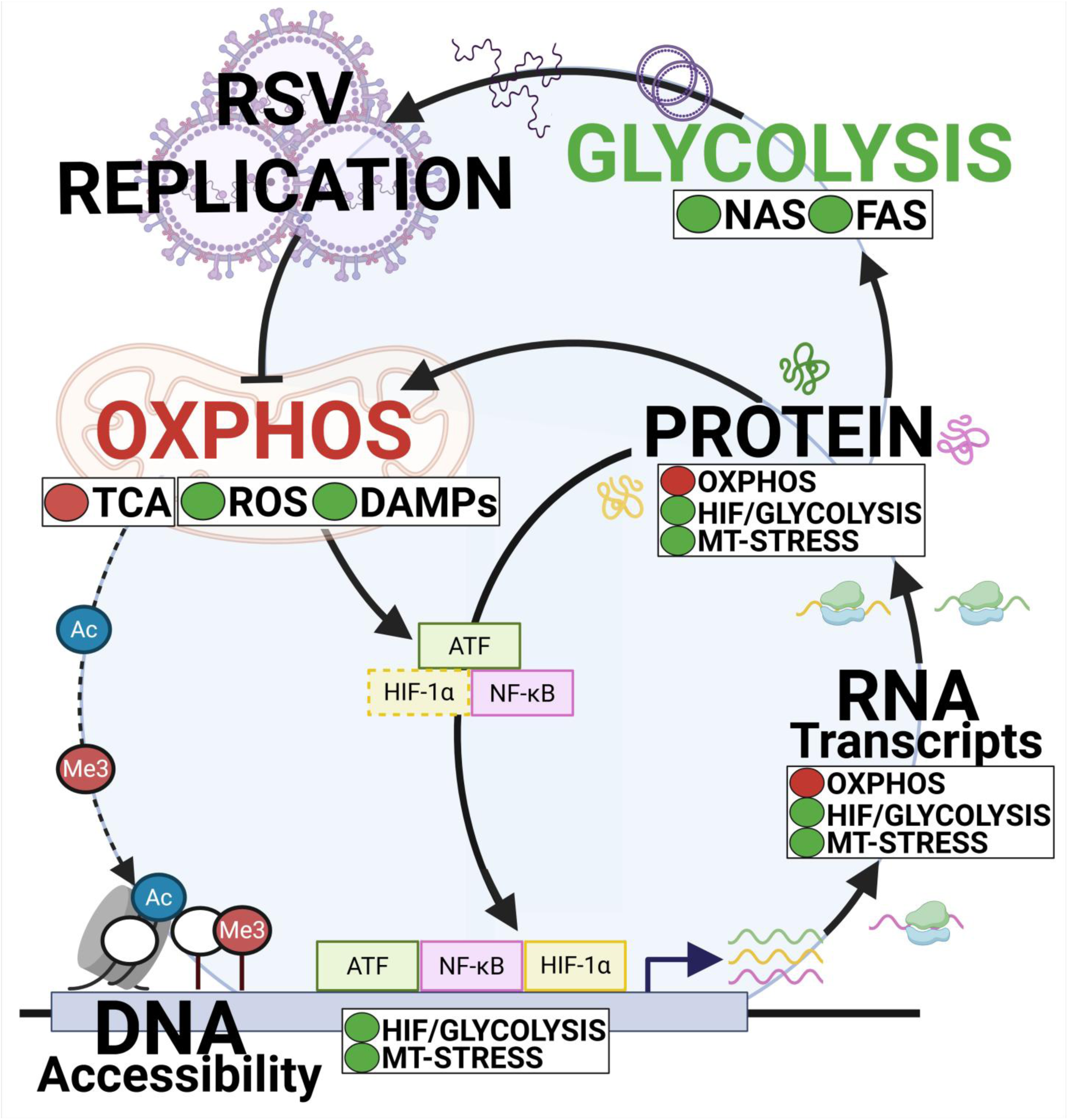
Proposed model of RSV-induced mitochondrial dysfunction across acute infection, severity, and long-term sequelae.

## DISCUSSION

Mitochondria function as essential signaling hubs that regulate cellular energetics, reactive oxygen species, calcium homeostasis, and the release of mitochondrial damage-associated molecular patterns^43–45^. Because of their central role in coordinating metabolism and immune signaling, viruses frequently hijack mitochondrial pathways through direct viral protein interactions, as well as transcriptional, epigenetic, and post-translational modifications that reprogram host cellular function^2,3,5,46–48^.

This phenomenon is increasingly recognized across severe respiratory viral infections^2,3,5,46,47^. The emergence of SARS-CoV-2 accelerated research into virus-host metabolic interactions and provided important mechanistic insights into how respiratory viruses exploit mitochondrial function. Our previous work demonstrated that SARS-CoV-2 rapidly reprograms host metabolism during acute infection^2,6–13,15,16^ through direct interference with mitochondrial respiratory complexes^2,7,10–12^, repression of OXPHOS transcripts^2,6,8,11^, and activation of HIF-1a and mTOR signaling pathways^6,8,13^.

During acute infection, SARS-CoV-2 suppresses OXPHOS, leading to increased mROS production that stabilizes HIF-1α. This drives a metabolic shift toward aerobic glycolysis and the pentose phosphate pathway, generating nucleotide and lipid precursors required for efficient viral replication. Concurrent activation of mTORC1 further reinforces this Warburg-like metabolic phenotype. In parallel, elevated mROS promotes the release of mtDAMPs, amplifying inflammatory signaling. Consistent with this model, inhibition of glycolysis, the pentose phosphate pathway, or HIF-1a/mTORC1 signaling reduces viral replication, whereas interventions that suppress mitochondrial function worsen disease. Conversely, mitochondria-targeted therapies that reduce oxidative stress mitigate disease severity across animal models, organoids, human monocytes, and cell culture systems, highlighting mitochondrial dysfunction as a central driver of SARS-CoV-2 pathogenesis^2,6–9,13^. Similar findings have been reported in IAV infection in mitochondrial complex-I deficient mice, where mitochondrial dysfunction is associated with impaired respiration, increased glycolysis, and worsened pathology^3^. Together, these studies identify mitochondrial dysfunction as a shared hallmark of respiratory viral infection.

Similarly to SARS-CoV-2 and IAV, RSV-induced mitochondrial dysfunction has been reported in multiple independent studies. RSV infection has been shown to impair OXPHOS, alter mitochondrial morphology, reduce ΔΨm^5,17,18^. Mechanistically, expression of the RSV Matrix (M) protein suppresses mitochondrial transcripts^17^, while RSV infection has also been shown to reduce NRF2 signaling through transcriptional and post-translational mechanisms, further exacerbating oxidative stress^19–22^. Consistent with this, complex I-deficient models exhibit increased viral replication and ROS production following RSV infection^49^. RSV infection further induces HIF-1α and its downstream transcriptional targets^23,24^, and pharmacologic inhibition of ROS or HIF-1α signaling reduces viral replication^23,25,26^.

Despite these observations, current evidence is derived from fragment experimental models, time points, and patient populations. As such, it remains unclear whether mitochondrial dysfunction represents a consistent and conserved feature of RSV infection across disease severity, viral strains, and clinical outcomes, or whether these metabolic alterations persist beyond acute infection to contribute to long-term sequelae.

To address this gap in RSV, we performed a comprehensive multi-omic analysis integrating 13 independent datasets spanning epigenomics, transcriptomics, proteomics, metabolomics, and single-cell sequencing across multiple model systems, viral strains, and clinical cohorts (**Supplemental Table 1**). Across these datasets, we identified a conserved RSV-associated metabolic signature characterized by suppression of OXPHOS, activation of HIF-1α signaling, and induction of mitochondrial stress pathways (**Summarized in Fig 7**). This was accompanied by metabolic remodeling that shifted cellular metabolism away from oxidative metabolism toward glycolysis, generating biosynthetic intermediates that may support viral replication. We further propose that impaired mitochondrial function increases mitochondrial stress and ROS production, promoting mtDAMP release into the cytosol and extracellular environment. These mtDAMPs can activate broad innate immune signaling pathways, potentially contributing to cytokine dysregulation and severe RSV pathology.

Consistent with this model, human adult challenge studies demonstrated progressive suppression of OXPHOS pathways during acute infection, with the most severe symptomatic patients exhibiting sustained OXPHOS suppression alongside elevated HIF-1α, mitochondrial stress, and immune activation signatures (**Fig 2**). In contrast, asymptomatic individuals demonstrated recovery of OXPHOS pathways and enrichment of multiciliated epithelial repair programs. Similar transcription patterns were observed in pediatric ICU cohorts, where severe disease was associated with greater metabolic dysfunction (**Fig 2**).

Importantly, our findings extend beyond acute infection, we identified long-term metabolic alterations following RSV infection (**Fig 4-5**). Single-cell analysis of nasal airway epithelial cells collected 1 YPI revealed persistent OXPHOS suppression, particularly in individuals who developed wheezing. These patients also exhibited depletion of multiciliated epithelial cells and expansion of secretory populations, suggesting persistent epithelial remodeling (**Fig 4**). Complementary blood methylation and serum metabolomic datasets further demonstrated sustained mitochondrial stress signatures and metabolic dysfunction long after viral clearance (**Fig 5**). These findings suggest that prolonged mitochondrial dysfunction may contribute to post-RSV complications in susceptible individuals.

We also identified strain-specific differences in metabolic reprogramming (**Fig 6**). While all RSV strains suppressed OXPHOS, RSV B:18537 exhibited weaker HIF-1α activation and reduced transcriptional disruption compared with other strains. Specifically, B:18537 enriched for mTOR signaling, suggesting it may utilize mTOR as a compensatory metabolic pathway due to an inability to fully stabilize HIF-1α, or alternatively, as a distinct, less-pathogenic survival strategy^50^. Notably, this strain has been associated with milder disease, suggesting that the magnitude of mitochondrial dysfunction may partially contribute to strain-dependent severity. Furthermore, while B:18537 is a historic lineage, ON1 and BA represent the dominant contemporary circulating lineages. These modern strains, characterized by large nucleotide duplications in the G gene, have largely swept the globe^28^. The observation that ON1 and BA exhibit more robust OXPHOS suppression and higher HIF-1α activation than historic strains underscores a potential evolutionary mechanism of viral adaptation. As viruses evolve under immune or therapeutic pressures, the efficiency with which they hijack host cellular machinery likely dictates their competitive fitness, thus, optimizing the host metabolic environment for viral replication appears to be a selectable trait. Future directions include Integrating these high-dimensional transcriptomic and metabolic signatures in a phylodynamic framework which will permit the reconstruction of how these host-response traits evolve concurrently with viral genetic diversity. Ultimately, such modeling is required to quantify whether enhanced metabolic hijacking acts as a primary driver of lineage fixation or a secondary consequence of genetic drift and shift.

Finally, we demonstrate that vaccination attenuates RSV-induced metabolic dysfunction (**Fig 3**). Vaccinated mice showed reduced metabolic disruption compared with unvaccinated animals, supporting the link between mitochondrial dysfunction and disease severity.

Collectively, these findings identify mitochondrial dysfunction as a central feature of RSV pathogenesis that spans acute infection severity, strain-specific responses, and long-term complications. The mechanistic parallels observed between RSV and SARS-CoV-2 suggest that mitochondrial hijacking may represent a conserved strategy used by respiratory viruses to enhance replication while simultaneously driving host pathology. These findings highlight mitochondrial-targeted therapeutics, antioxidant strategies, and modulation of HIF-1α signaling as promising approaches for reducing both acute disease severity and long-term complications following respiratory viral infection.

### Study Limitations

This study has several limitations. First, our analyses relied on publicly available datasets generated across independent studies with differences in sample collection, sequencing platforms, patient demographics, and infection timelines, which may introduce study-specific confounders. However, this heterogeneity also enabled identification of conserved signals across diverse systems. Second, our conclusions are based primarily on observational multi-omic analyses and pathway-level inference and therefore do not establish direct mechanistic causality between mitochondrial dysfunction and RSV severity or long-term outcomes. Third, several longitudinal pediatric cohorts, particularly the snRNA-seq dataset, had limited sample sizes, which may affect generalizability. Fourth, serum metabolomic datasets may not fully capture tissue-specific metabolic alterations within respiratory or immune compartments. Finally, although our findings suggest mitochondrial dysfunction may contribute to long-term complications such as wheezing, these datasets cannot fully distinguish RSV-driven effects from underlying host susceptibility factors. Additional mechanistic and prospective studies will be needed to validate these findings and assess whether mitochondrial-targeted interventions can improve RSV outcomes. Specifically, future studies should investigate whether the strain-specific metabolic differences observed i*n vitro* result in differential patterns in the airway epithelium, thereby directly linking viral evolutionary dynamics to the severity of long-term epithelial remodeling. This framework will be critical for evaluating transitioning viral dynamics and host phenotypic plasticity during the implementation of novel public health interventions, such as the widespread deployment of monoclonal antibodies like nirsevimab.

## Resource Availability

### Lead contact

Further information and requests for resources, data and code availability should be directed to the corresponding authors Joseph W. Guarnieri; Robert E. Schwartz; Nidia Sequeira Trovao.

### Materials availability

This study did not generate new, unique reagents or materials. All datasets analyzed in this research were obtained from publicly accessible datasets. The specific references and dataset accession numbers are listed in the “Experimental Model and Subject Details” below. Custom pathways used for gene set analysis are described in detail in “Methods Details” below.

### Data and code availability

All data used in this study are publicly available and can be accessed through their respective repository using the accession numbers listed in the “Experimental Model and Subject Details” section below. Any additional information required to reanalyze the data reported in this paper is available from the lead contact upon request.

### Experimental Model and Subject Details

#### Overview

This manuscript integrates data from a diverse set of experimental models. Each dataset was originally generated by independent investigators and is described in full methodological detail within the corresponding publications; readers are referred to those citations for comprehensive information.

In brief, data extracted from each source are summarized as follows RNA-seq datasets: R1(^30^); R2(^31^); R3(^31^); R4(^33^); R6(^34^); R7(^35^); R8(^35^); R9(^32^); snRNA-seq datasets: R5(^29^); Proteomics: P1(^38^). Metabolomics: M1(^39^); M2(^40^); ATAC-seq: A1(^30^); Bisulfide-sequencing: B1(^36^). Detailed experimental information for each dataset is provided in **Supplemental Table 1.** Below, we also provide a concise summary of the studies and sample types analyzed in this paper for comparative multi-omics integration.

#### RNA-seq, ATAC-seq, and Proteomics Analyses of In Vitro Infections Across Human Airway Models

##### RNA-seq (A549, hTBECs, NHBE-ALI)

Publicly available RNA-sequencing datasets were compiled to assess host transcriptional responses to RSV infection across multiple *in vitro* human airway models. Three independent datasets were included (R1-R3). The first dataset (R1(^30^); GEO:GSE161849) profiled RSV infection in A549 alveolar epithelial cells at 0.75 and 1 days post-infection (DPI), with matched infected (RSV+) and uninfected (RSV−) controls at each timepoint (n = 4 per group). The second dataset (R2(^31^); R3(^31^); GEO:GSE155152) examined A549 cells at 1, 2 and 4 DPI (1 DPI: RSV+ n = 3, RSV− n = 3; 2 DPI: RSV+ n = 2, RSV− n = 2; 4 DPI: RSV+ n = 3, RSV− n = 3). The same study also included human tracheobronchial epithelial cells (hTBECs) at 3 DPI (R3(^31^); GEO:GSE270463; RSV+ n = 3, RSV− n = 3). The third dataset (R4(^33^); GEO:GSE146795) evaluated primary normal human bronchial epithelial cells cultured at air-liquid interface (NHBE-ALI) at 6 DPI (RSV+ n = 4, RSV− n = 3). All three studies used RSV strain A2.

##### RNA-seq, Strain-Specific (A549)

Strain-specific transcriptional responses were assessed in A549 cells at 2.5 DPI, comparing RSV+ and RSV− conditions across four strains (R9(^32^); GEO:GSE196385). These 4 strains included two RSV-A strains, RSV/A/USA/BCM-Tracy/1989 (GA1) and RSV/A/USA/BCM813013/2013 (ON), and two RSV-B strains, RSV/B/WashingtonDC.USA/18537/1962 (GB1) and RSV/B/USA/BCM80171/2010 (BA), with matched controls for each strain (RSV+ n = 4, RSV− n = 4 per strain).

##### ATAC-seq (A549)

ATAC-seq data were analyzed to characterize RSV-induced chromatin accessibility changes in A549 cells. Cells were infected with RSV A2 and harvested at 1 DPI. Differential chromatin accessibility was assessed by comparing RSV-infected cells to uninfected controls (RSV+ n = 2, RSV− n = 2) (A1(^30^); GEO:GSE161849).

##### Proteomics (A549)

Proteomics data were analyzed to characterize RSV-driven changes in protein abundance in A549 cells, infected with RSV A2 and harvested at 1 DPI. Differential protein expression was assessed by comparing RSV-infected samples to uninfected controls (RSV+ n = 4, RSV− n = 4) (P1(^38^); ProteomeXchange: PXD004737).

#### RNA-seq and snRNA-seq Analyses of RSV Infection in Human Nasal Samples

##### RNA-seq (Adult Nasal Mucosa)

Nasal curettage samples from healthy adult volunteers (18-55 years) were collected at <1, 3, and 7 DPI following natural RSV infection, comparing RSV-infected individuals with symptoms (RSV+Symptoms+) and without symptoms (RSV+Symptoms−) to RSV-negative controls (RSV−). These data were published across two independent studies: one investigating transcriptomic changes in infected patients at <1 and 3 DPI (R6(^34^); GEO:GSE155237), at <1 DPI: RSV+Symptoms+ (n = 15) vs. RSV− (n = 14); RSV+Symptoms− (n = 15) vs. RSV− (n = 5); at 3 DPI: RSV+Symptoms+ (n = 5) vs. RSV− (n = 5), and a second study analyzing patients at <1 and 7 DPI (R7(^35^); GEO:GSE166161), at 7 DPI: RSV+Symptoms+ (n = 16) vs. RSV− (n = 16); RSV+Symptoms− (n = 16) vs. RSV− (n = 7). By analyzing data from both studies alongside their respective controls, we investigated progressive transcriptional changes across <1, 3, and 7 DPI.

##### RNA-seq (Infant Nasopharyngeal)

Nasopharyngeal samples from pediatric subjects (<2 years) with severe and mild RSV infection were collected 3 days post ICU (DP-ICU) admission. GSEA comparisons were performed comparing severe vs. mild 3 DP-ICU RSV infants (ICU-severe n = 27, ICU-mild n = 24) (R8(^35^); GEO:GSE146925).

##### snRNA-seq (Infant NAECs)

Primary NAECs were collected from children aged 2-3 years, at 1 YPI enrolled in the INSPIRE birth cohort and stratified by RSV infection status during infancy and wheeze phenotype. snRNA-seq was performed on differentiated NAEC cultures. Two comparisons were conducted: RSV-infected wheezers versus non-infected non-wheezers (RSV+Wheeze+ vs. RSV−Wheeze−; n = 2 per group) and RSV-infected non-wheezers versus non-infected non-wheezers (RSV+Wheeze− vs. RSV−Wheeze−; n = 2 per group) (R5(^29^); GEO:GSE286262).

#### Bisulfite Sequencing and Metabolomics Analyses of RSV-Associated Wheezing Outcomes in Infant Blood and Serum

##### Bisulfite Sequencing (Infant Whole Blood)

Genome-wide DNA methylation profiles were obtained from whole blood samples of pediatric subjects (<2 years) collected at 1 YPI following RSV infection (B1(^36^) GEO:GSE199334). Differential methylation analysis compared RSV-positive children who subsequently developed recurrent wheezing to those who did not (RSV+Wheeze+ vs. RSV+Wheeze−; n = 36 vs. n = 32).

##### Metabolomics (Infant Serum)

Infants ≤6 months of age hospitalized with a first episode of severe RSV bronchiolitis were enrolled and followed longitudinally through early childhood (up to ∼3 years) to examine serum metabolomic signatures associated with subsequent recurrent wheezing. Serum samples collected at follow-up visits 1-2.5 YPI were analyzed by untargeted ultra-performance liquid chromatography (UPLC) coupled with high-resolution tandem mass spectrometry (HR-MS/MS) metabolomics. Differential metabolite profiles were assessed by comparing RSV-positive infants who subsequently developed recurrent wheezing to those who did not (RSV+Wheeze+ vs. RSV+Wheeze−; n = 27 vs. n = 24) (M1(^39^)).

##### Metabolomics Analysis of RSV Infection and Vaccine Adjuvanting in a Murine Model

Metabolic alterations associated with RSV infection and vaccine adjuvanting were characterized in a BALB/c mouse model. Female BALB/c mice (6-8 weeks old) were intranasally immunized with a truncated RSV F protein (ΔF) formulated with the TriAdj adjuvant cocktail (poly(I:C), innate defense regulator peptide (IDR1002), and polyphosphazene (PCEP)), or left non-immunized, prior to RSV A2 challenge. Serum samples were collected at 7 DPI and subjected to untargeted metabolomic profiling. Two comparisons were performed: RSV-infected adjuvanted mice versus RSV-negative controls (RSV+TriAdj+ vs. RSV−; n = 3 per group) and RSV-infected non-adjuvanted mice versus RSV-negative controls (RSV+TriAdj− vs. RSV−; n = 3 per group) (M2(^40^)).

## Method Details

### Data Acquisition and Processing (RNA-seq, snRNA-seq)

All bulk RNA-seq [R1(^30^); R2(^31^); R3(^31^); R4(^33^); R6(^34^); R7(^35^); R8(^35^); R9(^32^)] and snRNA-seq R5(^29^) datasets were obtained from previously published studies, with library preparation, sequencing, and initial preprocessing described in the original sources (Experimental Model and Subject Details for more details). Bulk RNA-seq and scRNA-seq data were processed using standard alignment and quantification pipelines to generate raw gene-level count matrices^6,9,51–53^.

Bulk RNA-seq and scRNA-seq count tables were imported into DESeq2 (v1.46.0)^54^. Count matrices from all RNA-seq modalities were assembled into DESeqDataSet objects with defined experimental designs. After internal size-factor normalization, DESeq2 fit negative binomial generalized linear models and computed all pairwise contrasts, yielding normalized counts, log2 fold changes, standard errors, Wald statistics, p-values, and FDR-adjusted p-values. Gene annotations were appended using AnnotationDbi (v1.66.0), org.Hs.eg.db (v3.18.0), and online retrieval via biomaRt (v2.58.0) when needed.

Gene-level results were used for visualization, pathway scoring, and enrichment analyses. Volcano plots were generated with EnhancedVolcano (v1.14.0) (https://github.com/kevinblighe/EnhancedVolcano); heatmaps with ComplexHeatmap (v2.9.4)^55^, and circlize (v0.4.12); and lollipop plots using ggplot2 (v3.5.0) for plotting, ggh4x (v0.2.5) for facet_grid2 layouts, and scales (v1.3.0) for gradient and size transformations, with grDevices used internally for color handling.

### Metabolomic Data Processing and Differential Abundance Analysis

Targeted LC-MS/MS and flow injection analysis-tandem mass spectrometry (FIA-MS/MS) metabolomic datasets were preprocessed and normalized according to their source publications M1 (^39^); and M2(^40^). Differential metabolite abundance was analyzed using limma (v3.52.4)^56^, with a design matrix constructed from sample metadata and linear models fitted for each metabolite. Empirical Bayes moderation, with trend and robust options, was applied to stabilize variance estimates. Pairwise contrasts were computed to identify alterations in key metabolic pathways, including energy metabolism, amino acid and nucleotide metabolism, lipid metabolism, and redox balance. Resulting log₂ fold changes, t-statistics, p-values, and false discovery rate (FDR)-adjusted p-values were compiled, and Volcano plots were generated with EnhancedVolcano (v1.14.0) (https://github.com/kevinblighe/EnhancedVolcano) to visualize metabolite-level changes.

### Proteomic Sample Preparation and Differential Analysis

Protein abundance was quantified using the SomaScan aptamer platform, measuring over ∼6,000 serum proteins according to their source publications P1(^38^). Expression matrices were generated from raw seq.* identifiers, and a limma (v3.52.4)^56^. Linear modeling framework was applied with design matrices derived from groupings (RSV-infected, mock-infected). Empirical Bayes moderation stabilized variance estimates, and all pairwise contrasts were computed to identify protein-level changes associated with Seq.* identifiers were mapped to gene symbols and UniProt IDs, then cross-referenced to official Human Genome Organisation (HUGO) Gene Nomenclature Committee (HGNC) symbols using biomaRt (v2.58.0) for downstream protein-level analysis and visualization. Volcano plots were generated with EnhancedVolcano (v1.14.0) (https://github.com/kevinblighe/EnhancedVolcano); heatmaps with ComplexHeatmap (v2.9.4)^55^ and circlize (v0.4.12).

### ATAC-seq Differential Expression Analysis

Data was obtained from previously published studies A1(^30^), and initial preprocessing was performed as previously described (^30^). After which Peak-level count matrices were analyzed in R using DESeq2 (v1.46.0) (doi:10.1186/s13059-014-0550-8). After normalization, negative binomial generalized linear models were fitted to identify differentially accessible regions between experimental groups. Pairwise contrasts were computed to generate log_₂_ fold changes, statistical significance, and FDR-adjusted p-values. Peaks were annotated to nearby genes to enable gene-level interpretation. Downstream pathway and gene set enrichment analyses were performed using the same frameworks applied to RNA-seq data.

### Bisulfide-seq Differential Expression Analysis

Bisulfite sequencing datasets were obtained from previously published studies, with experimental procedures and preprocessing described in the original sources B1(^34^). Methylation data were analyzed in R using limma (v3.52.4)^56^, with appropriate transformation of methylation values where required. Linear models were fitted for each cytosine-phosphate-guanine (CpG) site or region, followed by empirical Bayes moderation to assess differential methylation between groups. Pairwise contrasts were used to generate effect sizes, p-values, and FDR-adjusted p-values. Differentially methylated sites were annotated to nearby genes for downstream interpretation. Pathway and gene set enrichment analyses were performed using the same approaches applied to RNA-seq data.

### Custom Pathways

To examine extracellular matrix (ECM) remodeling, we evaluated genes associated with the *Extracellular Matrix Organization* pathway as defined in the Reactome database (Reactome ID: R-HSA-1474244). For senescence-related genes, we curated “Senescence,” “Induced Senescence,” and “Inhibited Senescence-associated genes” were compiled from the Database of Cell Senescence Genes (https://genomics.senescence.info/cells/).

All remaining pathways used in this study were derived from previously published custom metabolic and immune gene sets (^6,9^), with a few minor changes. For the metabolic pathways (^6^) related to mitochondrial biology, we incorporated the custom OXPHOS-focused gene lists and consolidated all mitochondrial OXPHOS-related pathways, including “Complex I-V,” “Complex I,” “Complex II,” “Complex III,” “Complex IV,” “MT-Ribosome,” “MT-Biogenesis,” “MT-Protein Import,” and “MT-CoQ Synthesis”, into a newly defined macro-category termed “OXPHOS.” For glycolytic and hypoxia-responsive pathways, we created an additional macro-category, “HIF/mTOR,” that integrates the “HIF1A,” “mTOR,” and “Glycolysis” pathways. Finally, from the custom immune pathway set (^9^), we established two broader categories to capture integrated stress and cell-death responses: “ISR & UPR,” which merges the integrated stress response and unfolded protein response pathways, and “PANoptosis,” which unifies the “Pyroptosis”, “Necroptosis”, and “Apoptosis” programs into a single composite pathway.

### Figure Preparation

Summary schematics and conceptual illustrations were created using BioRender (BioRender.com). BioRender was used exclusively for graphical representation and did not involve data manipulation or quantitative processing. Final figure layouts were assembled using standard vector graphic software.

#### Quantification and Statistical Analysis

All statistical analyses were performed in R (v4.3.0) using modality-appropriate packages. For transcriptomic analyses, DESeq2 (v1.46.0) was used for normalization and differential expression testing, with gene annotations obtained from AnnotationDbi (v1.66.0), org.Hs.eg.db (v3.18.0), and online retrieval via biomaRt (v2.58.0) when needed. Proteomic and metabolomic differential abundance analyses were performed using limma (v3.52.4)^56^. Unless otherwise specified, p-values were adjusted using the Benjamini-Hochberg method. For pathway-level analyses, an adjusted FDR < 0.25 threshold was applied for significance across all GSEA and gene set enrichment outputs. For visualizations displaying individual metabolites, proteins, or transcripts, including volcano plots, lollipop plots, and heatmaps, features meeting a threshold of p < 0.05 were considered significant.

Figures and data visualizations were generated using EnhancedVolcano (v1.14.0) (https://github.com/kevinblighe/EnhancedVolcano) for volcano plots, ComplexHeatmap (v2.9.4)^55^ and circlize (v0.4.12) for heatmaps, ggplot2 (v3.5.0) for plotting, ggh4x (v0.2.5) for advanced facet layouts, and scales (v1.3.0) for gradient and size transformations. Additional data handling and I/O packages included readxl and openxlsx for Excel import/export, and reshape2 for data reshaping.

## Author Contributions

Conceptualization: J.W.G., N.S.T., R.E.S; Methodology: J.W.G; Formal Analysis: J.W.G; Investigation: J.W.G., N.S.T., R.E.S.; Data Curation: J.W.G; Resources: J.W.G., N.S.T., R.E.S; Writing - Original Draft: J.W.G., N.S.T.; Writing - Review & Editing: J.W.G., N.S.T., R.E.S; Visualization: J.W.G.; Funding Acquisition: J.W.G., N.S.T., R.E.S; Supervision: J.W.G., N.S.T., R.E.S.

---

**Supplementary Figure 1.**
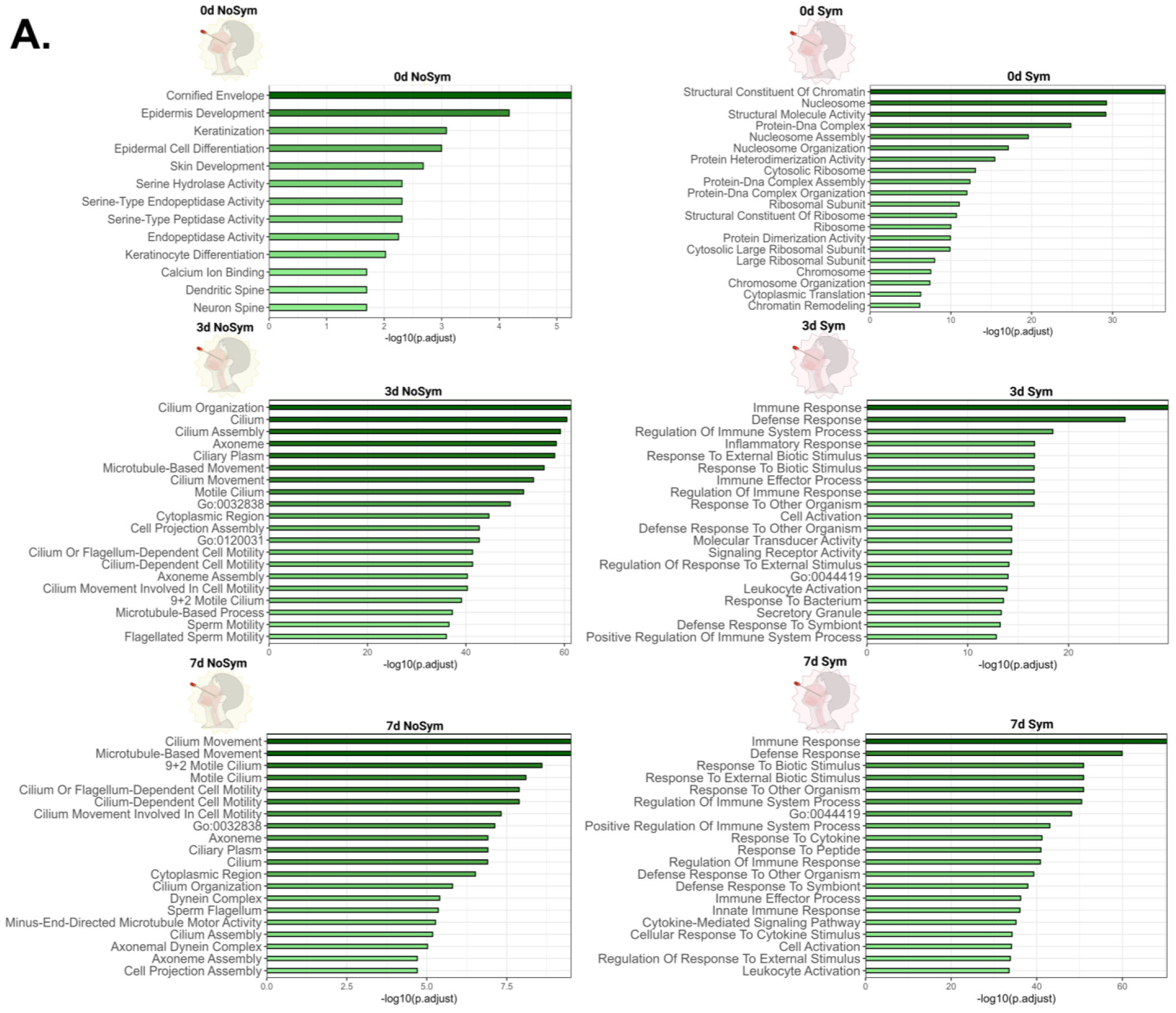
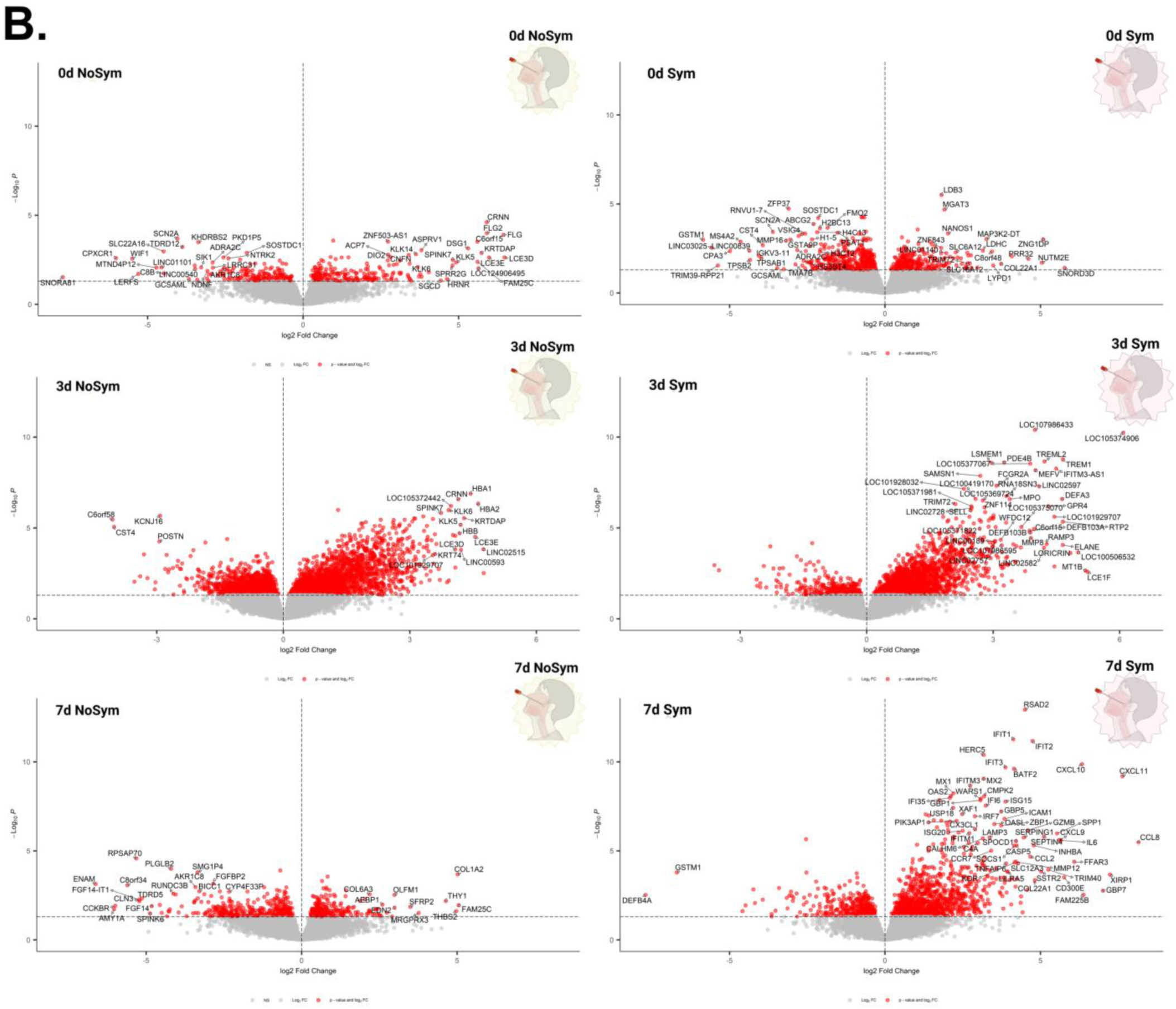
Symptomatic adult RSV infection is associated with sustained inflammatory signaling and delayed epithelial recovery. (A) GO enrichment analysis and (B) Volcano plots of from adult nasal curettage, comparisons include symptomatic (RSV+Symptoms+) and asymptomatic (RSV+Symptoms−) individuals relative to RSV-negative controls (RSV−) across <1, 3, and 7 DPI (R6(^32^); R7(^33^)). (B) Each point represents a transcript plotted by log_₂_ fold change (x-axis) versus −log_₁₀_ adjusted p-value (y-axis).

**Supplementary Figure 2.**
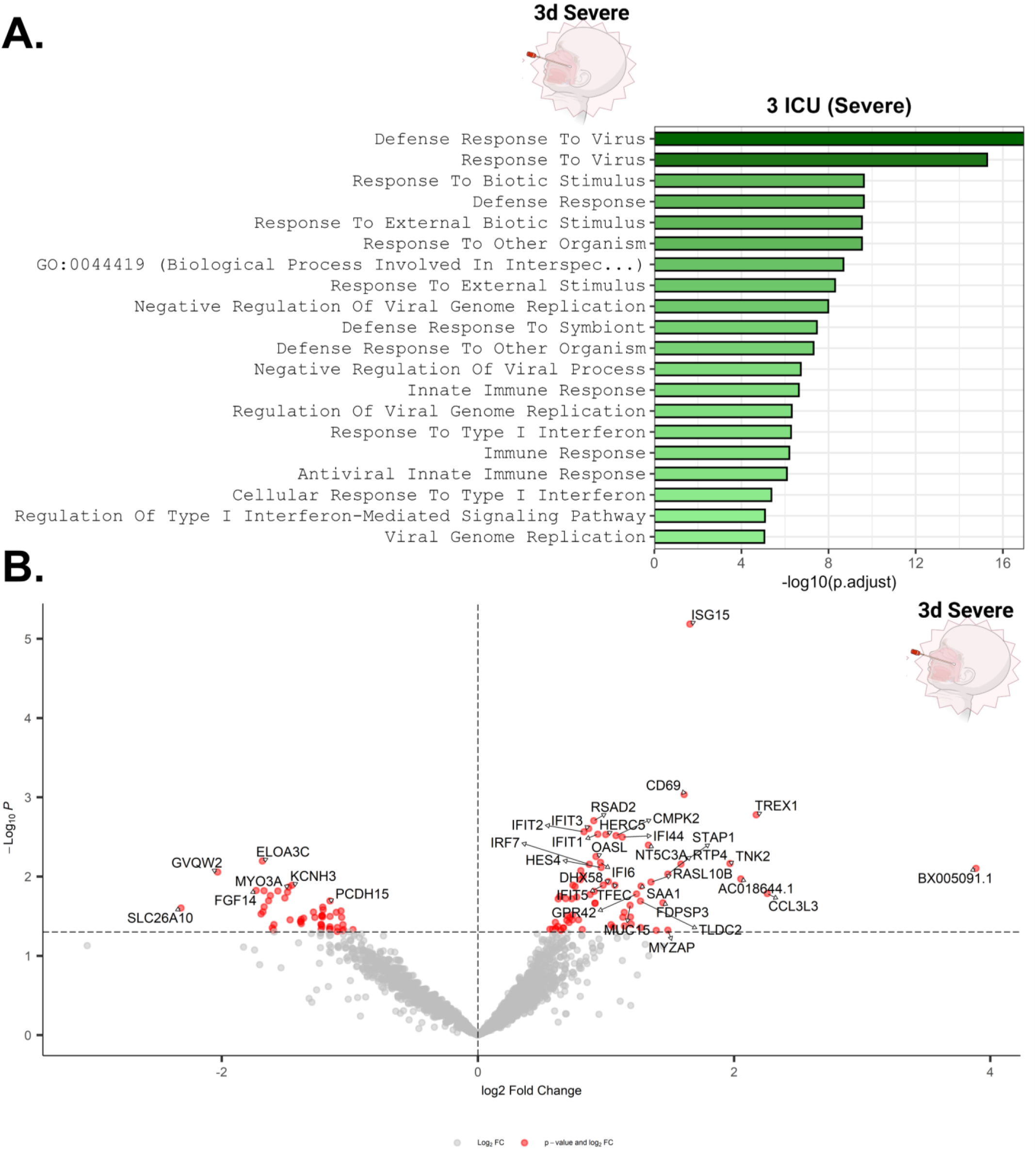
Severe pediatric RSV infection is associated with sustained inflammatory signaling and broader transcriptional dysregulation. (A) GO enrichment analysis and (B) volcano plots from pediatric nasopharyngeal RNA-seq comparisons between severe and mild RSV cases collected at 3 DP-ICU (R8(^33^)). (B) Each point represents a transcript plotted by log_₂_ fold change (x-axis) versus −log adjusted p-value (y-axis).

**Supplementary Figure 3.**
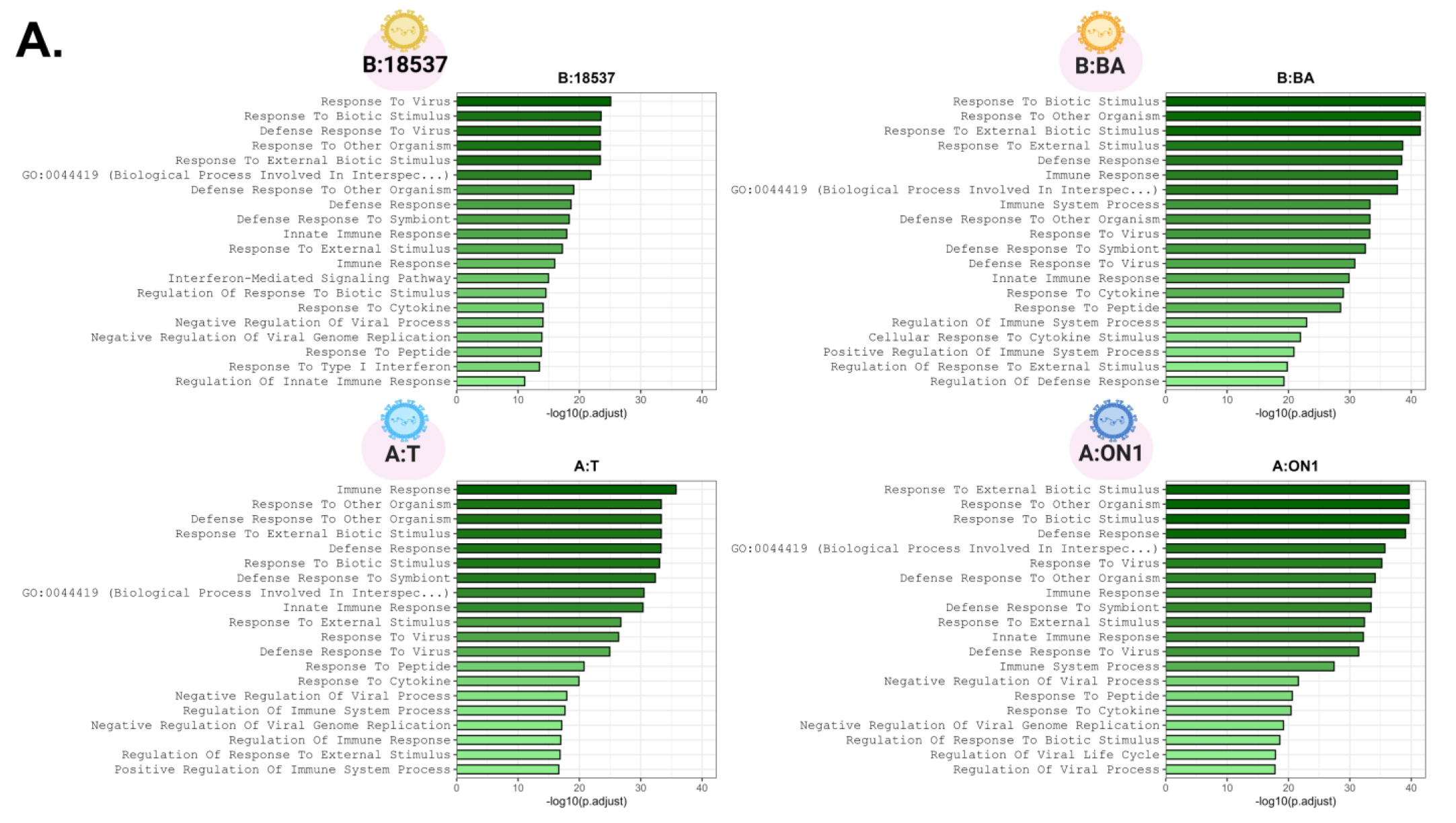
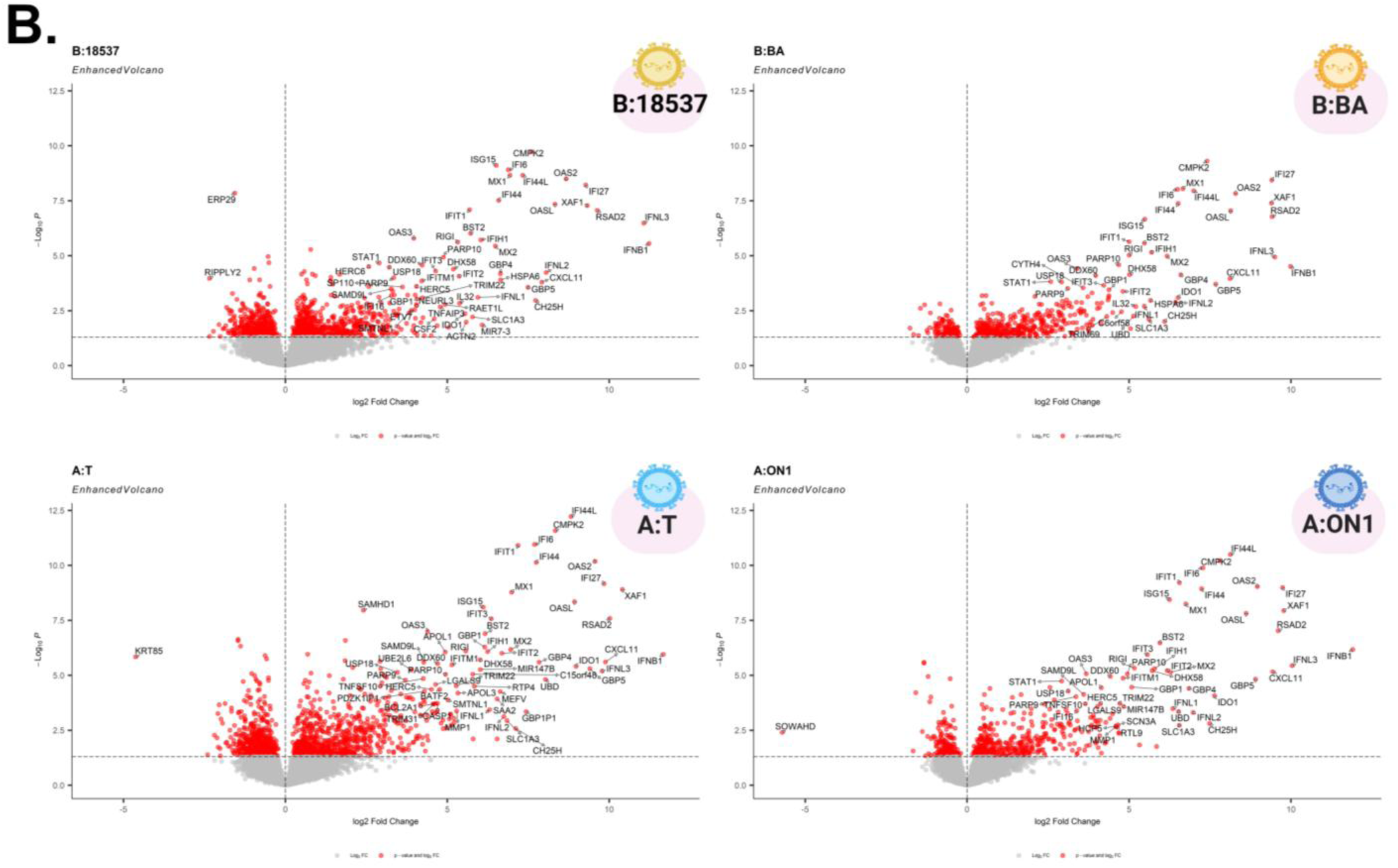
RSV strains exhibit distinct transcriptional and metabolic pathway responses. GO enrichment analysis and (B) volcano plots from RNA-seq comparisons across four RSV strains: A/Tracy (GA1), B/18537 (GB1), A/ON1 (ON), and B/BA (BA) relative to matched RSV-negative controls (R9(^32^)). (B) Each point represents a transcript plotted by log₂ fold change (x-axis) versus −log₁₀ adjusted p-value (y-axis).

**Supplemental Table 1.**
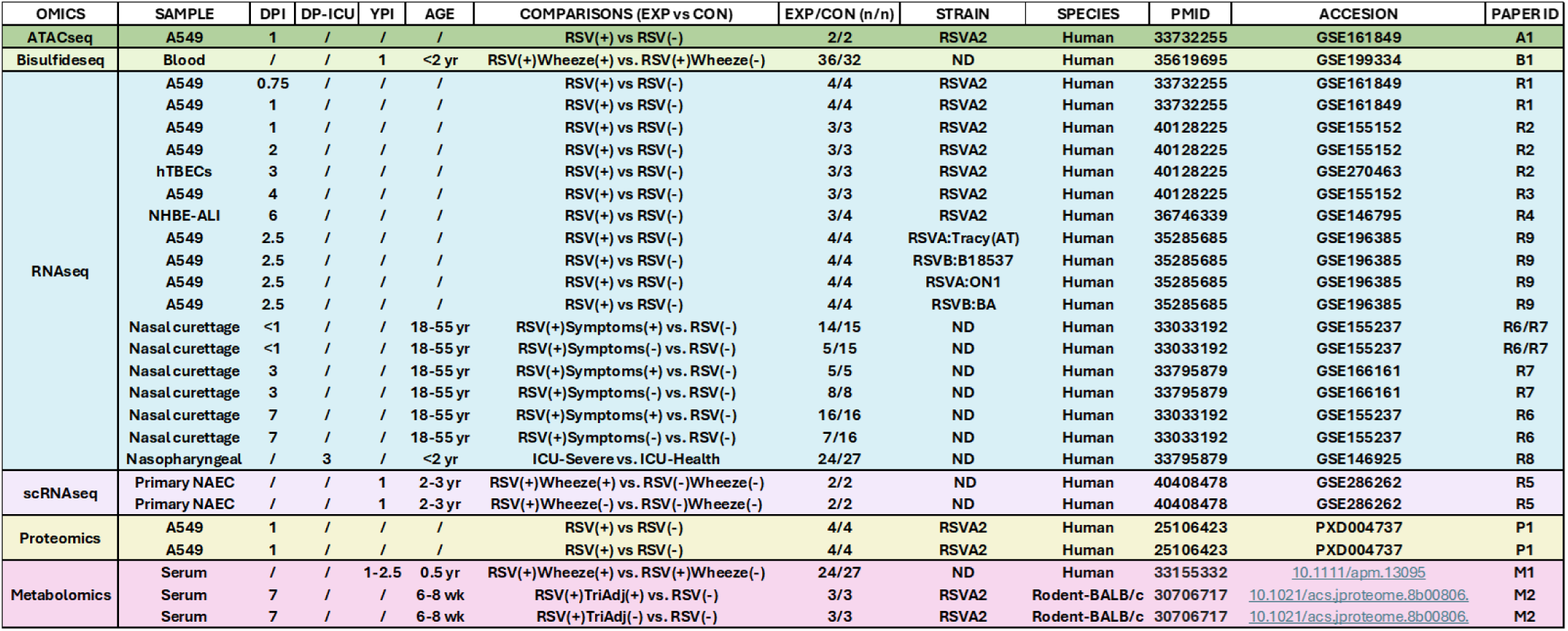
Summary of multi-omic datasets used for meta-analysis of RSV infection across human airway models.

## REFERENCES

1. Borchers, A. T., Chang, C., Gershwin, M. E. & Gershwin, L. J. Respiratory syncytial virus--a comprehensive review. Clin. Rev. Allergy Immunol. 45, 331–379 (2013).

2. Guarnieri, J. W. et al. SARS-CoV-2 mitochondrial metabolic and epigenomic reprogramming in COVID-19. Pharmacol. Res. 204, 107170 (2024).

3. Fuchs, A. L. et al. Mitochondrial dysfunction enhances influenza pathogenesis by up-regulating de novo sialic acid biosynthesis. Sci. Adv. 11, eadu3739 (2025).

4. Trovao, N. S. et al. Viral evolution in the cosmos. in Fundamentals of Space Medicine and Clinical Technology 67–96 (Elsevier, 2026). doi:10.1016/B978-0-443-32904-3.00009-1.

5. Hu, M. et al. Respiratory syncytial virus co-opts host mitochondrial function to favour infectious virus production. eLife 8, e42448 (2019).

6. Guarnieri, J. W. et al. Core mitochondrial genes are down-regulated during SARS-CoV-2 infection of rodent and human hosts. Sci. Transl. Med. 15, eabq1533 (2023).

7. Guarnieri, J. W. et al. SARS-COV-2 viroporins activate the NLRP3-inflammasome by the mitochondrial permeability transition pore. Front. Immunol. 14, 1064293 (2023).

8. Guarnieri, J. W. et al. Mitochondrial antioxidants abate SARS-COV-2 pathology in mice. Proc. Natl. Acad. Sci. U. S. A. 121, e2321972121 (2024).

9. Topper, M. J. et al. Lethal COVID-19 associates with RAAS-induced inflammation for multiple organ damage including mediastinal lymph nodes. Proc. Natl. Acad. Sci. U. S. A. 121, e2401968121 (2024).

10. Guarnieri, J. W. et al. TARGETED DOWN REGULATION OF CORE MITOCHONDRIAL GENES DURING SARS-COV-2 INFECTION. BioRxiv Prepr. Serv. Biol. 2022.02.19.481089 (2022) doi:10.1101/2022.02.19.481089.

11. Haltom, J. A. et al. Importance of De Novo Gene Evolution to Emerging Viral Threats: The ORF10 Strain-Restricted Orphan Gene of SARS-CoV-2 Promotes Pathogenesis. Mol. Biol. Evol. 42, msaf211 (2025).

12. McDonald, J. T., et al. Role of miR-2392 in driving SARS-CoV-2 infection. Cell Rep. 37, 109839 (2021).

13. Duan, X. et al. An airway organoid-based screen identifies a role for the HIF1α-glycolysis axis in SARS-CoV-2 infection. Cell Rep. 37, 109920 (2021).

14. Codo, A. C. et al. Elevated Glucose Levels Favor SARS-CoV-2 Infection and Monocyte Response through a HIF-1α/Glycolysis-Dependent Axis. Cell Metab. 32, 437–446.e5 (2020).

15. Jamison, D. A. et al. A comprehensive SARS-CoV-2 and COVID-19 review, Part 1: Intracellular overdrive for SARS-CoV-2 infection. Eur. J. Hum. Genet. EJHG 30, 889–898 (2022).

16. Narayanan, S. A. et al. A comprehensive SARS-CoV-2 and COVID-19 review, Part 2: host extracellular to systemic effects of SARS-CoV-2 infection. Eur. J. Hum. Genet. EJHG 32, 10–20 (2024).

17. Hu, M., Bogoyevitch, M. A. & Jans, D. A. Respiratory Syncytial Virus Matrix Protein Is Sufficient and Necessary to Remodel Host Mitochondria in Infection. Cells 12, 1311 (2023).

18. Elesela, S. et al. Sirtuin 1 regulates mitochondrial function and immune homeostasis in respiratory syncytial virus infected dendritic cells. PLoS Pathog. 16, e1008319 (2020).

19. Hosakote, Y. M., Liu, T., Castro, S. M., Garofalo, R. P. & Casola, A. Respiratory syncytial virus induces oxidative stress by modulating antioxidant enzymes. Am. J. Respir. Cell Mol. Biol. 41, 348–357 (2009).

20. Cho, H.-Y. et al. Murine Neonatal Oxidant Lung Injury: NRF2-Dependent Predisposition to Adulthood Respiratory Viral Infection and Protection by Maternal Antioxidant. Antioxidants 10, 1874 (2021).

21. Komaravelli, N. et al. Respiratory syncytial virus infection down-regulates antioxidant enzyme expression by triggering deacetylation-proteasomal degradation of Nrf2. Free Radic. Biol. Med. 88, 391–403 (2015).

22. Hosakote, Y. M. et al. Viral-mediated inhibition of antioxidant enzymes contributes to the pathogenesis of severe respiratory syncytial virus bronchiolitis. Am. J. Respir. Crit. Care Med. 183, 1550–1560 (2011).

23. Morris, D. R., Qu, Y., Agrawal, A., Garofalo, R. P. & Casola, A. HIF-1α Modulates Core Metabolism and Virus Replication in Primary Airway Epithelial Cells Infected with Respiratory Syncytial Virus. Viruses 12, 1088 (2020).

24. Haeberle, H. A. et al. Oxygen-independent stabilization of hypoxia inducible factor (HIF)-1 during RSV infection. PloS One 3, e3352 (2008).

25. Hosakote, Y. M. et al. Antioxidant mimetics modulate oxidative stress and cellular signaling in airway epithelial cells infected with respiratory syncytial virus. Am. J. Physiol. Lung Cell. Mol. Physiol. 303, L991–1000 (2012).

26. Mata, M., Morcillo, E., Gimeno, C. & Cortijo, J. N-acetyl-L-cysteine (NAC) inhibit mucin synthesis and pro-inflammatory mediators in alveolar type II epithelial cells infected with influenza virus A and B and with respiratory syncytial virus (RSV). Biochem. Pharmacol. 82, 548–555 (2011).

27. Langedijk, A. C. et al. The genomic evolutionary dynamics and global circulation patterns of respiratory syncytial virus. Nat. Commun. 15, 3083 (2024).

28. Yu, J.-M., Fu, Y.-H., Peng, X.-L., Zheng, Y.-P. & He, J.-S. Genetic diversity and molecular evolution of human respiratory syncytial virus A and B. Sci. Rep. 11, 12941 (2021).

29. Berdnikovs, S. et al. Single-cell profiling demonstrates the combined effect of wheeze phenotype and infant viral infection on airway epithelial development. Sci. Adv. 11, eadr9995 (2025).

30. Xu, X., et al. The SWI/SNF-Related, Matrix Associated, Actin-Dependent Regulator of Chromatin A4 Core Complex Represses Respiratory Syncytial Virus-Induced Syncytia Formation and Subepithelial Myofibroblast Transition. Front. Immunol. 12, 633654 (2021).

31. Kalita, P. et al. Molecular basis for human respiratory syncytial virus transcriptional regulator NS1 interactions with MED25. Nat. Commun. 16, 2883 (2025).

32. Rajan, A. et al. Multiple Respiratory Syncytial Virus (RSV) Strains Infecting HEp-2 and A549 Cells Reveal Cell Line-Dependent Differences in Resistance to RSV Infection. J. Virol. 96, e0190421 (2022).

33. Talukdar, S. N. et al. RSV-induced expanded ciliated cells contribute to bronchial wall thickening. Virus Res. 327, 199060 (2023).

34 Habibi, M. S. et al. Neutrophilic inflammation in the respiratory mucosa predisposes to RSV infection. Science 370, eaba9301 (2020).

35. Felt, S. A. et al. Detection of respiratory syncytial virus defective genomes in nasal secretions is associated with distinct clinical outcomes. Nat. Microbiol. 6, 672–681 (2021).

36. Pischedda, S. et al. Role and Diagnostic Performance of Host Epigenome in Respiratory Morbidity after RSV Infection: The EPIRESVi Study. Front. Immunol. 13, 875691 (2022).

37. Munday, D. C. et al. Quantitative proteomic analysis of A549 cells infected with human respiratory syncytial virus. Mol. Cell. Proteomics MCP 9, 2438–2459 (2010).

38. Dave, K. A. et al. A comprehensive proteomic view of responses of A549 type II alveolar epithelial cells to human respiratory syncytial virus infection. Mol. Cell. Proteomics MCP 13, 3250–3269 (2014).

39. Zhang, X. et al. Serum metabolomic profiling reveals important difference between infants with and without subsequent recurrent wheezing in later childhood after RSV bronchiolitis. APMIS Acta Pathol. Microbiol. Immunol. Scand. 129, 128–137 (2021).

40. Sarkar, I., Zardini Buzatto, A., Garg, R., Li, L. & van Drunen Littel-van den Hurk, S. Metabolomic and Immunological Profiling of Respiratory Syncytial Virus Infection after Intranasal Immunization with a Subunit Vaccine Candidate. J. Proteome Res. 18, 1145–1161 (2019).

41. Topol, E. J. COVID-19 can affect the heart. Science 370, 408–409 (2020).

42. Cai, M., Xie, Y., Topol, E. J. & Al-Aly, Z. Three-year outcomes of post-acute sequelae of COVID-19. Nat. Med. 30, 1564–1573 (2024).

43. Stojanovic, L. et al. ZNFX1 Functions as a Master Regulator of Epigenetically Induced Pathogen Mimicry and Inflammasome Signaling in Cancer. Cancer Res. 85, 1183–1198 (2025).

44. Tasoula, A., Poignant, F., Guarnieri, J. W., Schwertz, H. & Nelson, G. A. Health impacts of radiation in space and countermeasures. in Fundamentals of Space Medicine and Clinical Technology 467–488 (Elsevier, 2026). doi:10.1016/B978-0-443-32904-3.00034-0.

45. Zhu, Y. et al. PET Imaging with [18F]ROStrace Detects Oxidative Stress and Predicts Parkinson’s Disease Progression in Mice. Antioxidants 13, 1226 (2024).

46. Muccilli, S. G. et al. Mitochondrial hyperactivity and reactive oxygen species drive innate immunity to the yellow fever virus-17D live-attenuated vaccine. PLoS Pathog. 21, e1012561 (2025).

47. Chia, S. B. et al. Respiratory viral infections awaken metastatic breast cancer cells in lungs. Nature 645, 496–506 (2025).

48. Davanzo, G. G. et al. Obesity-Induced Metabolic Priming Exacerbates SARS-CoV-2 Inflammation. Immunology 175, 323–338 (2025).

49. Hu, M., Bogoyevitch, M. A. & Jans, D. A. Subversion of Host Cell Mitochondria by RSV to Favor Virus Production is Dependent on Inhibition of Mitochondrial Complex I and ROS Generation. Cells 8, 1417 (2019).

50. Cheng, S.-C. et al. mTOR- and HIF-1α-mediated aerobic glycolysis as metabolic basis for trained immunity. Science 345, 1250684 (2014).

51. Houerbi, N. et al. Secretome profiling reveals acute changes in oxidative stress, brain homeostasis, and coagulation following short-duration spaceflight. Nat. Commun. 15, 4862 (2024).

52. Guarnieri, J. W. et al. Methodologies for Mitochondrial Omic Profiling During Spaceflight. Methods Mol. Biol. 2878, 273–291 (2025).

53. McDonald, J. T. et al. Space radiation damage rescued by inhibition of key spaceflight associated miRNAs. Nat. Commun. 15, 4825 (2024).

54. Love, M. I., Huber, W. & Anders, S. Moderated estimation of fold change and dispersion for RNA-seq data with DESeq2. Genome Biol. 15, 550 (2014).

55. Gu, Z., Eils, R. & Schlesner, M. Complex heatmaps reveal patterns and correlations in multidimensional genomic data. Bioinformatics 32, 2847–2849 (2016).

56. Ritchie, M. E. et al. limma powers differential expression analyses for RNA-sequencing and microarray studies. Nucleic Acids Res. 43, e47 (2015).

