## Supplementary Materials for "Multi-omic analysis identifies mitochondrial dysfunction as a conserved driver of acute severity and long-term complications in RSV infection"

| OMICS | SAMPLE | DPI | DP-ICU | YPI | AGE | COMPARISONS (EXP vs CON) | EXP/CON (n/n) | STRAIN | SPECIES | PMID | ACCESION | PAPER ID |
|---|---|---|---|---|---|---|---|---|---|---|---|---|
| ATACseq | A549 | 1 | / | / | / | RSV(+) vs RSV(-) | 2/2 | RSVA2 | Human | 33732255 | GSE161849 | A1 |
| Bisulfideseq | Blood | / | / | 1 | <2 yr | RSV(+)Wheeze(+) vs. RSV(+)Wheeze(-) | 36/32 | ND | Human | 35619695 | GSE199334 | B1 |
| RNAseq | A549 | 0.75 | / | / | / | RSV(+) vs RSV(-) | 4/4 | RSVA2 | Human | 33732255 | GSE161849 | R1 |
| | A549 | 1 | / | / | / | RSV(+) vs RSV(-) | 4/4 | RSVA2 | Human | 33732255 | GSE161849 | R1 |
| | A549 | 1 | / | / | / | RSV(+) vs RSV(-) | 3/3 | RSVA2 | Human | 40128225 | GSE155152 | R2 |
| | A549 | 2 | / | / | / | RSV(+) vs RSV(-) | 3/3 | RSVA2 | Human | 40128225 | GSE155152 | R2 |
| | hTBECs | 3 | / | / | / | RSV(+) vs RSV(-) | 3/3 | RSVA2 | Human | 40128225 | GSE270463 | R2 |
| | A549 | 4 | / | / | / | RSV(+) vs RSV(-) | 3/3 | RSVA2 | Human | 40128225 | GSE155152 | R3 |
| | NHBE-ALI | 6 | / | / | / | RSV(+) vs RSV(-) | 3/4 | RSVA2 | Human | 36746339 | GSE146795 | R4 |
| | A549 | 2.5 | / | / | / | RSV(+) vs RSV(-) | 4/4 | RSVA:Tracy(AT) | Human | 35285685 | GSE196385 | R9 |
| | A549 | 2.5 | / | / | / | RSV(+) vs RSV(-) | 4/4 | RSVB:B18537 | Human | 35285685 | GSE196385 | R9 |
| | A549 | 2.5 | / | / | / | RSV(+) vs RSV(-) | 4/4 | RSVA:ON1 | Human | 35285685 | GSE196385 | R9 |
| | A549 | 2.5 | / | / | / | RSV(+) vs RSV(-) | 4/4 | RSVB:BA | Human | 35285685 | GSE196385 | R9 |
| | Nasal curettage | <1 | / | / | 18-55 yr | RSV(+)Symptoms(+) vs. RSV(-) | 14/15 | ND | Human | 33033192 | GSE155237 | R6/R7 |
| | Nasal curettage | <1 | / | / | 18-55 yr | RSV(+)Symptoms(-) vs. RSV(-) | 5/15 | ND | Human | 33033192 | GSE155237 | R6/R7 |
| | Nasal curettage | 3 | / | / | 18-55 yr | RSV(+)Symptoms(+) vs. RSV(-) | 5/5 | ND | Human | 33795879 | GSE166161 | R7 |
| | Nasal curettage | 3 | / | / | 18-55 yr | RSV(+)Symptoms(-) vs. RSV(-) | 8/8 | ND | Human | 33795879 | GSE166161 | R7 |
| | Nasal curettage | 7 | / | / | 18-55 yr | RSV(+)Symptoms(+) vs. RSV(-) | 16/16 | ND | Human | 33033192 | GSE155237 | R6 |
| | Nasal curettage | 7 | / | / | 18-55 yr | RSV(+)Symptoms(-) vs. RSV(-) | 7/16 | ND | Human | 33033192 | GSE155237 | R6 |
| | Nasopharyngeal | / | 3 | / | <2 yr | ICU-Severe vs. ICU-Health | 24/27 | ND | Human | 33795879 | GSE146925 | R8 |
| scRNAseq | Primary NAEC | / | / | 1 | 2-3 yr | RSV(+)Wheeze(+) vs. RSV(-)Wheeze(-) | 2/2 | ND | Human | 40408478 | GSE286262 | R5 |
| | Primary NAEC | / | / | 1 | 2-3 yr | RSV(+)Wheeze(-) vs. RSV(-)Wheeze(-) | 2/2 | ND | Human | 40408478 | GSE286262 | R5 |
| Proteomics | A549 | 1 | / | / | / | RSV(+) vs RSV(-) | 4/4 | RSVA2 | Human | 25106423 | PXD004737 | P1 |
| | A549 | 1 | / | / | / | RSV(+) vs RSV(-) | 4/4 | RSVA2 | Human | 25106423 | PXD004737 | P1 |
| Metabolomics | Serum | / | / | 1-2.5 | 0.5 yr | RSV(+)Wheeze(+) vs. RSV(+)Wheeze(-) | 24/27 | ND | Human | 33155332 | 10.1111/apm.13095 | M1 |
| | Serum | 7 | / | / | 6-8 wk | RSV(+)TriAdj(+) vs. RSV(-) | 3/3 | RSVA2 | Rodent-BALB/c | 30706717 | 10.1021/acs.jproteome.8b00806. | M2 |
| | Serum | 7 | / | / | 6-8 wk | RSV(+)TriAdj(-) vs. RSV(-) | 3/3 | RSVA2 | Rodent-BALB/c | 30706717 | 10.1021/acs.jproteome.8b00806. | M2 |

**Supplemental Table 1. Summary of multi-omic datasets used for meta-analysis of RSV infection across human airway models.**

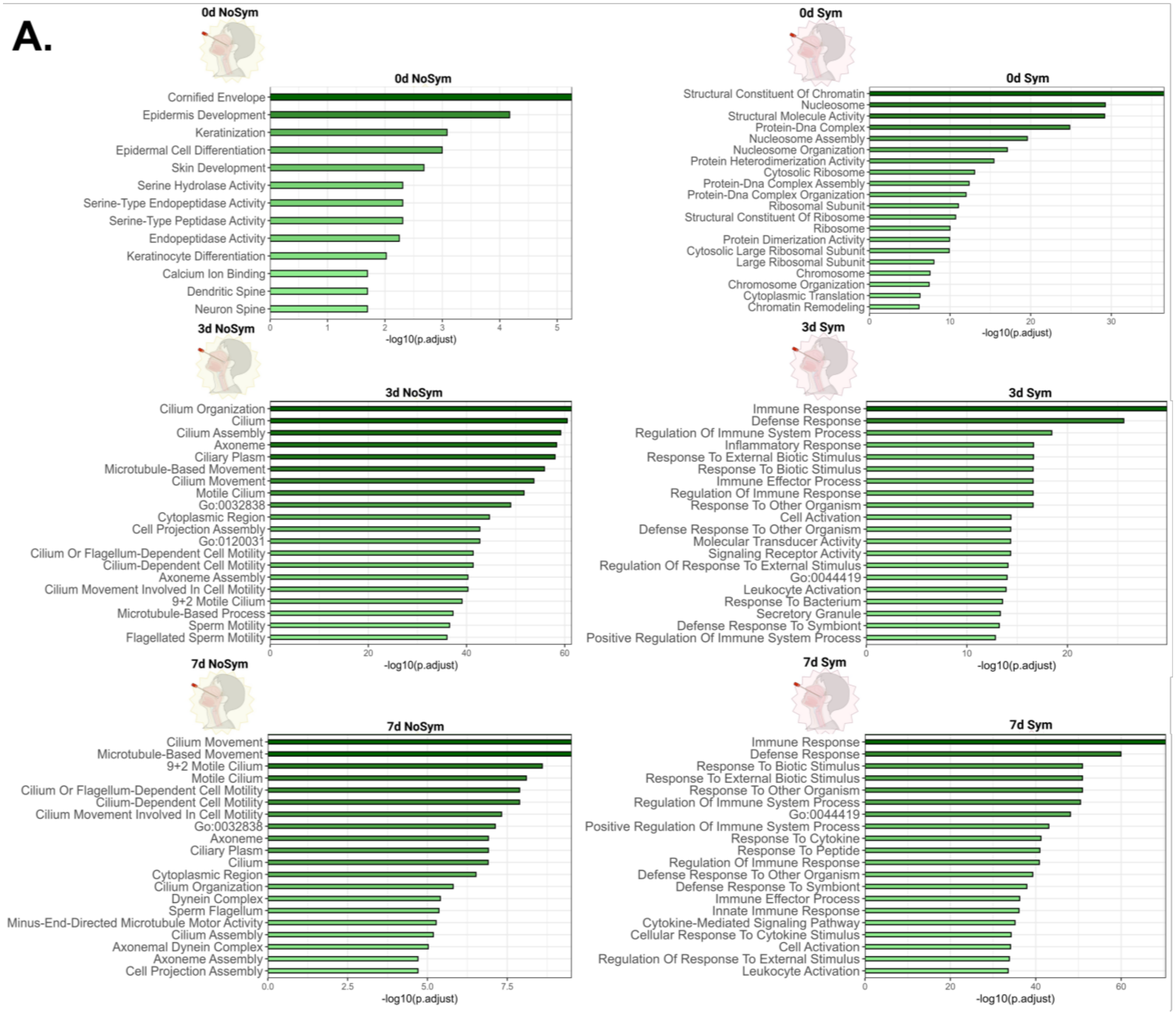


**Supplementary Figure 1. Symptomatic adult RSV infection is associated with sustained inflammatory signaling and delayed epithelial recovery.** (A) GO enrichment analysis and (B) Volcano plots of from adult nasal curettage, comparisons include symptomatic (RSV+Symptoms+) and asymptomatic (RSV+Symptoms−) individuals relative to RSV-negative controls (RSV−) across <1, 3, and 7 DPI (R6([32]); R7([33])). (B) Each point represents a transcript plotted by $\log_2$ fold change (x-axis) versus $-\log_{10}$ adjusted p-value (y-axis).

**B.**

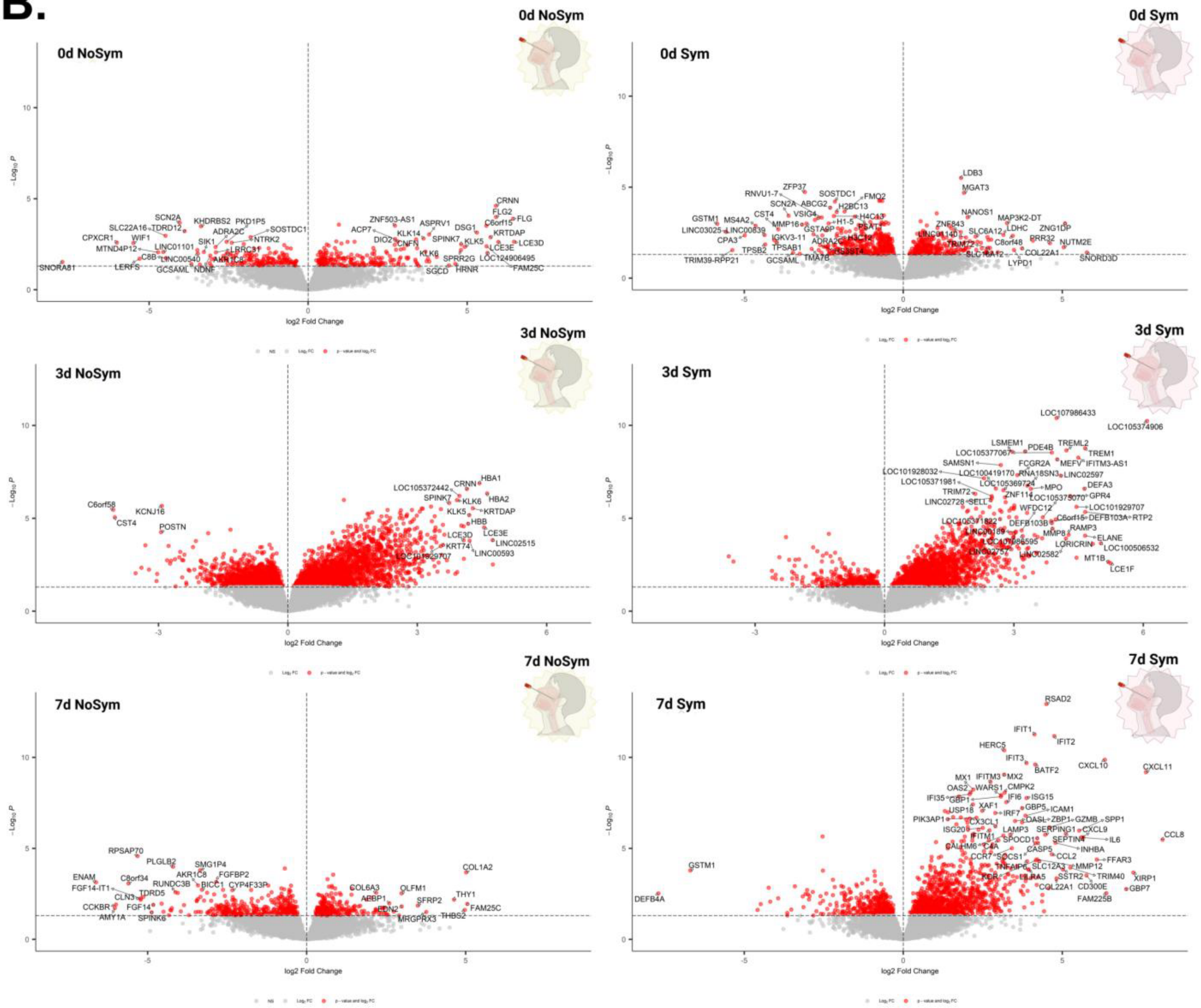


**Supplementary Figure 1. Symptomatic adult RSV infection is associated with sustained inflammatory signaling and delayed epithelial recovery (continued).**

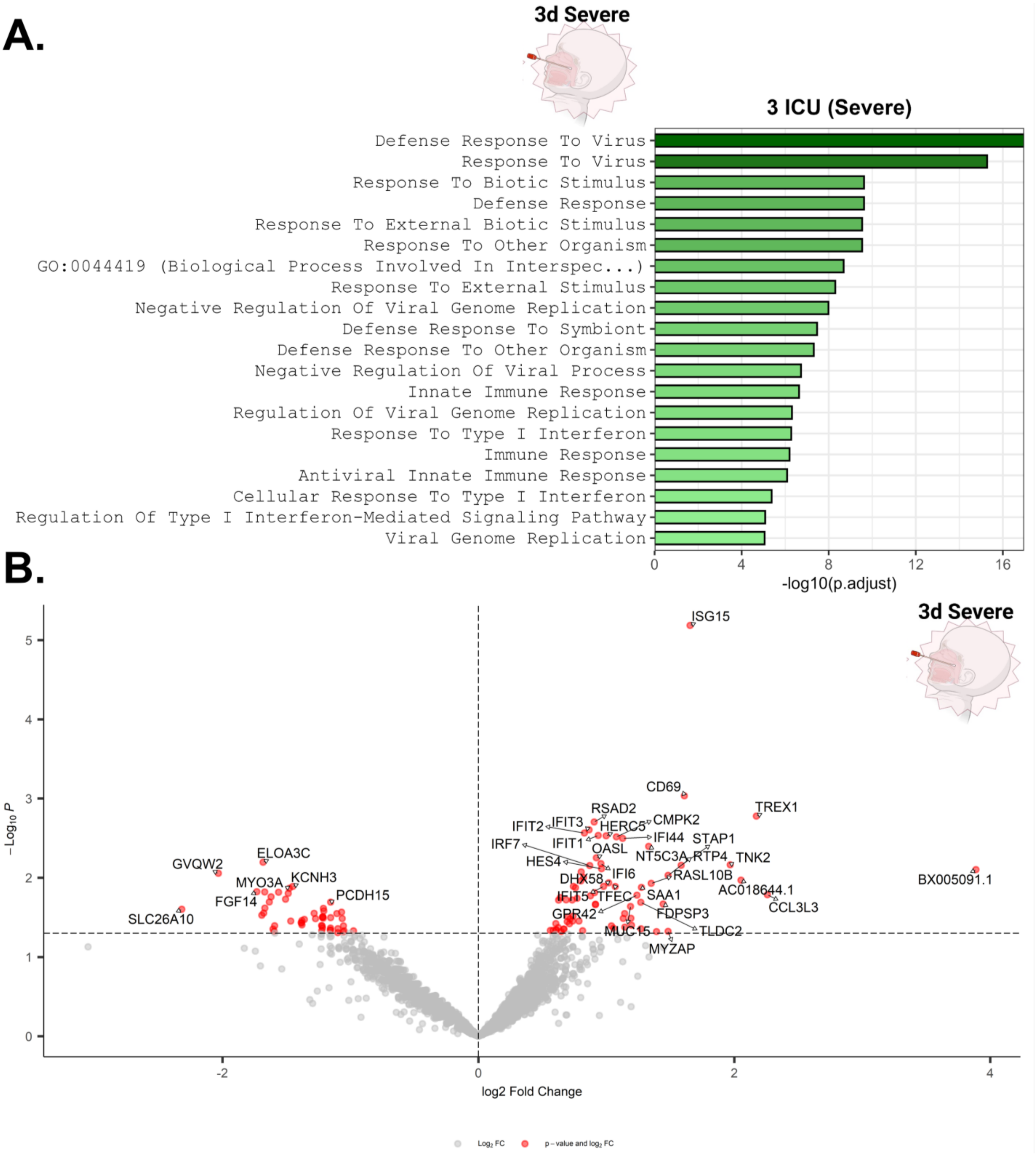


**Supplementary Figure 2. Severe pediatric RSV infection is associated with sustained inflammatory signaling and broader transcriptional dysregulation.** (A) GO enrichment analysis and (B) volcano plots from pediatric nasopharyngeal RNA-seq comparisons between severe and mild RSV cases collected at 3 DP-ICU (R8([33])). (B) Each point represents a transcript plotted by $\log_2$ fold change (x-axis) versus $-\log_{10}$ adjusted p-value (y-axis).

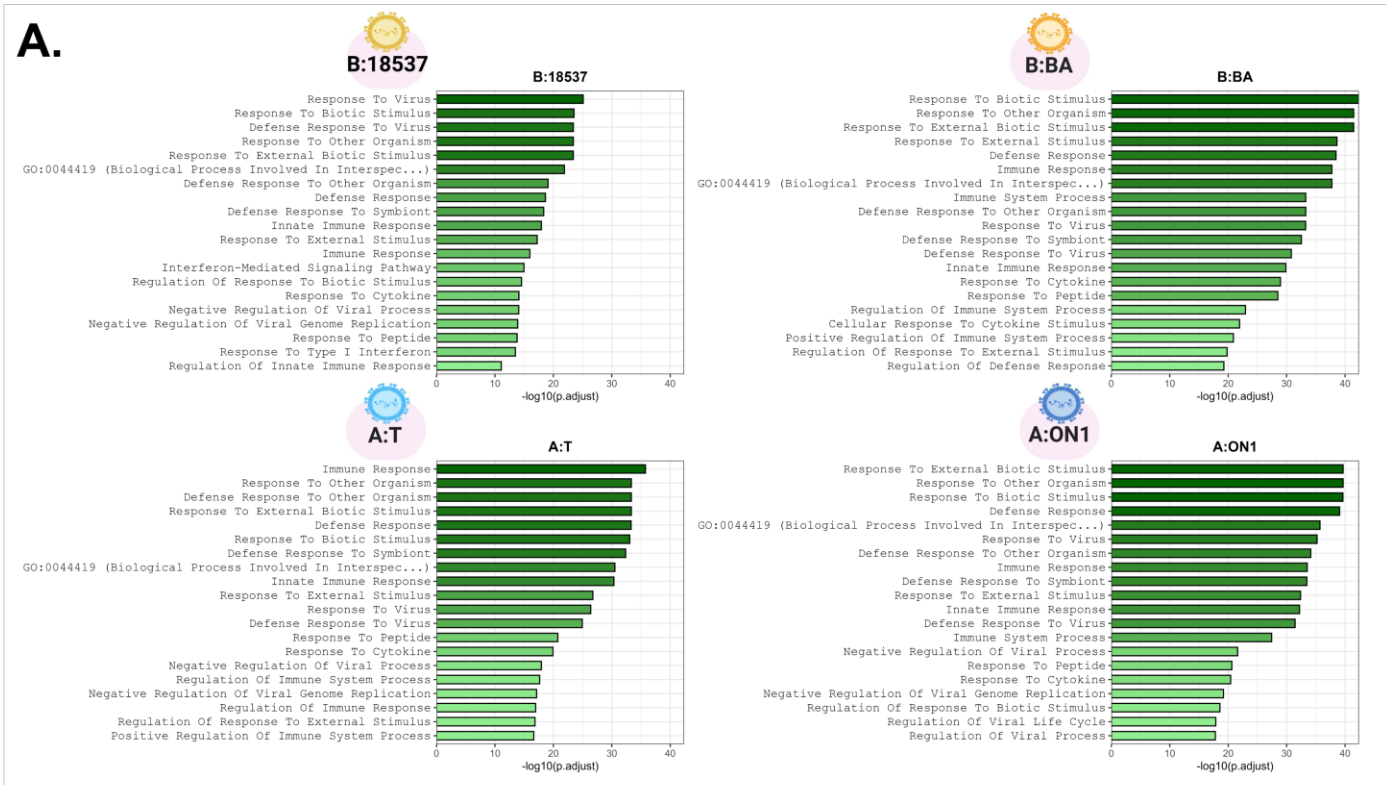


**Supplementary Figure 3. RSV strains exhibit distinct transcriptional and metabolic pathway responses.** GO enrichment analysis and (B) volcano plots from RNA-seq comparisons across four RSV strains: A/Tracy (GA1), B/18537 (GB1), A/ON1 (ON), and B/BA (BA) relative to matched RSV-negative controls (R9([32])). (B) Each point represents a transcript plotted by $\log_2$ fold change (x-axis) versus $-\log_{10}$ adjusted p-value (y-axis).

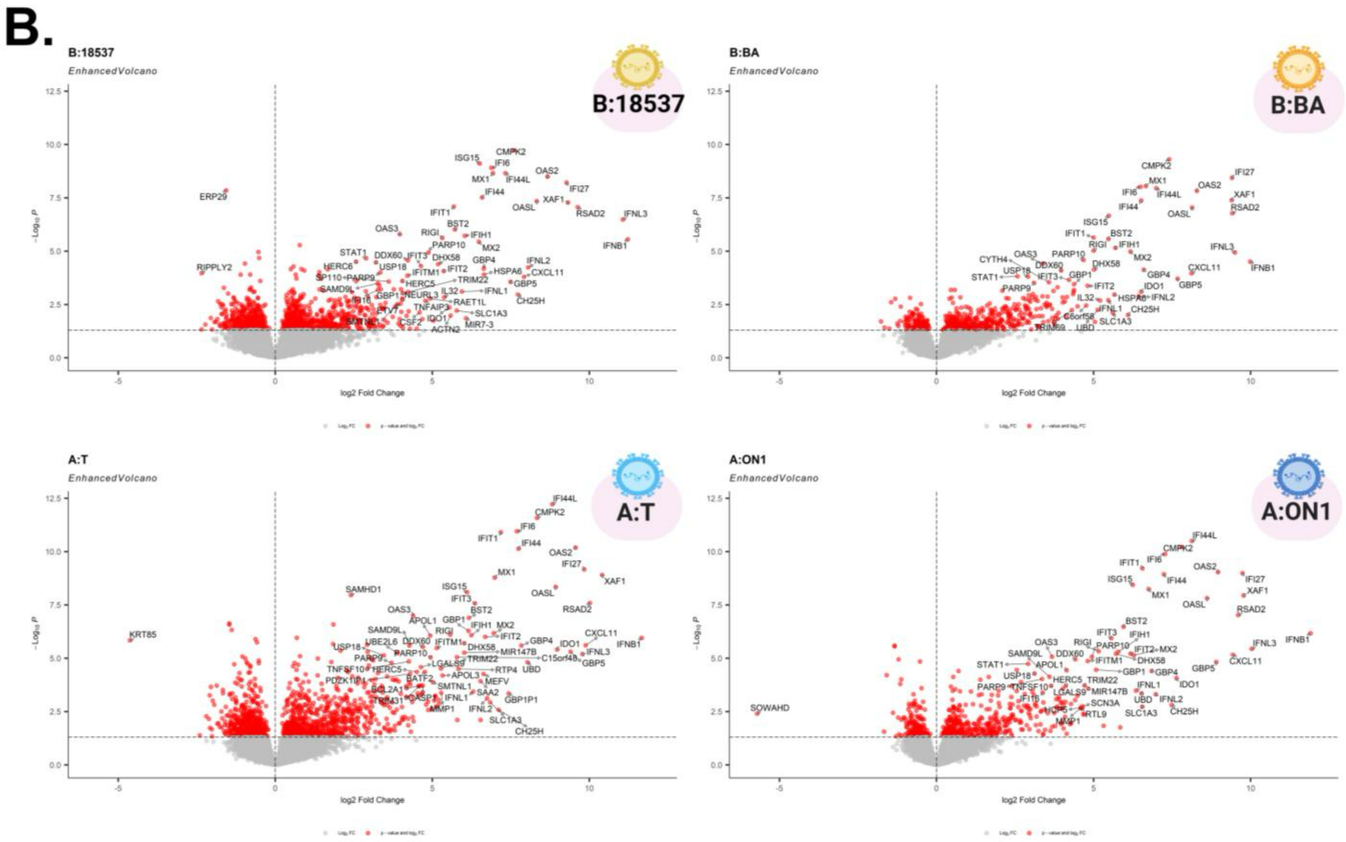


**Supplementary Figure 3. RSV strains exhibit distinct transcriptional and metabolic pathway responses (continued).**
